# Gaps in archaeological metadata reporting: a meta-analysis of human paleogenomic studies in Western Eurasia

**DOI:** 10.1101/2025.07.16.665041

**Authors:** Robert Staniuk, Victor Yan Kin Lee, Adrian Timpson, Kristian Kristiansen, Stephen Shennan, Fernando Racimo, Mark G. Thomas

## Abstract

Paleogenomic research has dramatically increased our understanding of past demographic and adaptive processes, but has also been criticized for a perceived disconnect between geneticists and other parties involved in the study of the past. For interdisciplinary research to be productive, contextual metadata associated with paleogenomic samples should be accessible in the same publication. Here, we conduct a pilot study examining the extent of archaeological metadata reporting in 30 peer-reviewed human paleogenomic studies, based on genetic sequences from 3911 ancient humans predominately from Western Eurasia, published between 2013 and 2024. We show inconsistent reporting of archaeological data across studies, and have sought to identify the driving factors. Overall, we found no strong explanatory variables, though some metadata fields – like *Geolocation* – have improved in completeness over time. These inconsistencies mean that metadata reporting practice is often insufficient for assessing archaeological interpretations independently, and hinders potential investigations of the relationship between patterns of cultural and genetic changes using already published data. We further propose a checklist with the aim of initiating community discussions on developing a better practice for collecting and reporting archaeological data alongside human paleogenomic studies in the future.

## Introduction

The last 15 years of human paleogenomic research in archaeology have revolutionized our understanding of the past. Improvements in ancient genome extraction and sequencing, as well as new bioinformatic and inferential methods, have led to a dramatic increase in ancient DNA data and findings from around the world. This has allowed researchers to study biological relatedness (Mittnik et al. 2019; Popli et al. 2023; Cassidy et al. 2025), identify signatures of natural selection (Mathieson et al. 2015; Evershed et al. 2022; Barrie et al. 2024; Irving-Pease et al. 2024), gain insight into past demographic processes (Bramanti et al. 2009; Sarkissian et al. 2013; Lazaridis et al. 2014; Skoglund et al. 2014; Haak et al. 2015; Brace et al. 2019; Lazaridis et al. 2022; Allentoft, Sikora, Fischer, et al. 2024), and track domestication processes (Librado et al. 2024 June 6) in the past. This rapid pace of development has been facilitated by the public data archiving requirements that are de rigueur in the field of genetics, and provide researchers with open access to previously generated ancient DNA (aDNA) sequences (Ehler et al. 2019; Mallick et al. 2024; Schmid et al. 2024). Although ancient DNA analysis as a methodological approach has been welcomed by many in the archaeological community, criticisms have been raised about the way it has been deployed, interpreted, and presented (Hofmann 2015; Vander Linden 2016; Heyd 2017; Ion 2017; Johannsen et al. 2017; Furholt 2018; Booth 2019; Frieman and Hofmann 2019; Furholt 2019; Sykes et al. 2019). The issues raised include, but are not limited to, inferred relationships between genetically derived clusters and archaeological cultures (Heyd 2017; Ion 2017; Johannsen et al. 2017; Furholt 2018; Frieman and Hofmann 2019; Furholt 2019), sampling biases (Hofmann 2015; Vander Linden 2016; Ion 2017; Sykes et al. 2019), epistemological incongruence between the questions posed and processes inferred (Hofmann 2015; Ion 2017; Furholt 2019; Sykes et al. 2019), and a tendency to generalize rather than provide nuanced (Healy 2017) interpretations of findings (Hofmann 2015; Vander Linden 2016; Johannsen et al. 2017; Frieman and Hofmann 2019; Furholt 2019).

The more recent focus of human paleogenomic research on site-specific investigations signals the emergence of a more collaborative framework suitable for incorporating high-resolution archaeological and genetic data (Mittnik et al. 2019; Schroeder et al. 2019; Cassidy et al. 2020; Sjögren et al. 2020; Brami et al. 2022), and suggests macroscale studies can benefit from explicitly incorporating high-resolution cultural data (Racimo, Sikora, et al. 2020; Kristiansen 2022; Shennan 2024). If successful, this approach would stimulate research into the mechanisms responsible for the long-term patterns of human evolutionary change documented in archaeology (Perreault 2019; Kristiansen 2022; Shennan 2024). However, achieving this goal requires datasets incorporating aDNA sequences and detailed cultural, contextual and environmental information associated with the sampled individuals. Presently, only one of these datasets is widely available – aDNA – as archaeology continues to struggle with generating unified open-access datasets with standardized information on excavated human remains (Anagnostou et al. 2015; Reiter et al. 2024). This is precisely where archaeology needs to draw from the data archiving developments of DNA research, like INSDC (Karsch-Mizrachi et al. 2025), and propose not only a general standard of reporting archaeological data but also encourage its adoption into a database, where this information becomes recorded and easily available to other researchers adhering to the FAIR principles (Wilkinson et al. 2016; Reiter et al. 2024).

While a repository or database for reporting archaeological data of comparable scope and widespread utilization has yet to exist, archaeological data about sequenced individuals are routinely reported as metadata in the Supplementary Information (SI) of human paleogenomic studies, particularly those covering a broad geographical scale where individual cultural information is not directly relevant. Generalized cultural data are provided in the descriptive supplementary, while additional information such as geolocation or radiocarbon dating is typically published in a tabulated form. Together they represent the most up-to-date information on the spatiotemporal positioning of cultural traits linked to genomic data, and the most accessible complementary archaeological dataset for the published genomic data. However, even brief examination of the contents of relevant SI files indicates that reporting standards for archaeological metadata vary within a single study, as well as across different papers. These inconsistencies are well-known to the community working with human aDNA but the general consensus is that any data is better than none.

As available human aDNA sequences have increased exponentially in numbers (Mallick et al. 2024; Schmid et al. 2024) and interdisciplinary research on links between culture and genetics gains momentum, accessing quality archaeological data becomes crucial. Before we determine what infrastructure is needed to store specific data, it is necessary to assess metadata availability and its reporting standard. It is by evaluating the *status quo* that gaps and key variables can be identified and structural solutions can be developed.

To do this, we collated archaeological metadata reported in 30 human paleogenomic peer-reviewed articles published between 2013 and 2024 (Table 1). As a pilot study, we focused on macroscale studies with individuals sampled from Western Eurasia since the region has been the most intensively sampled in the field to date (Mallick et al. 2024), and unlike site-specific studies raw archaeological data are often overlooked. We recorded which of the 16 most archaeologically relevant metadata fields were available, and proposed 5 plausible explanatory factors (Table 2). We then explored variation across metadata and publications, looked for improvements in the availability of metadata over time, and tested how well the explanatory variables explained metadata availability.

**Table 1:**
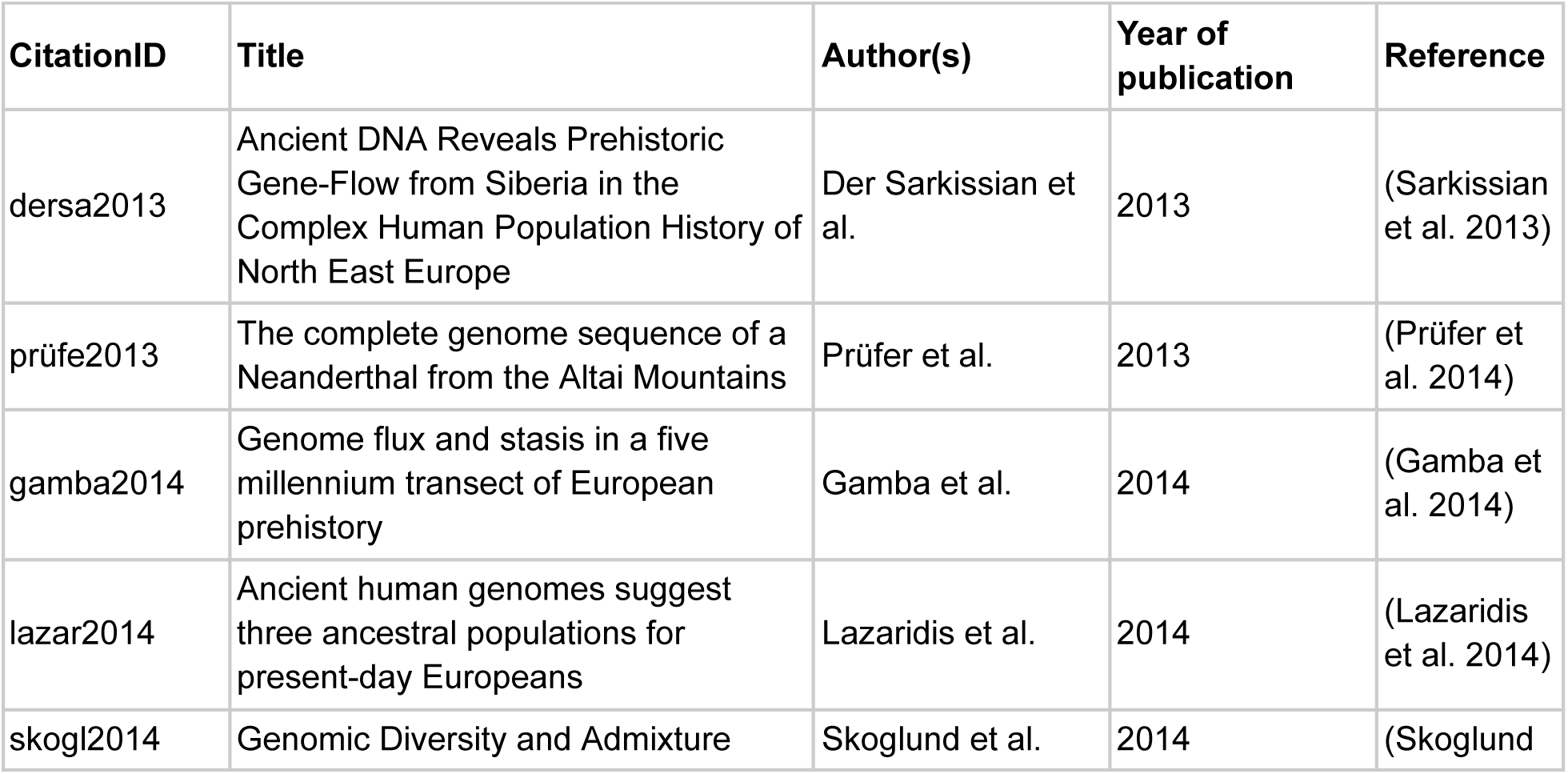
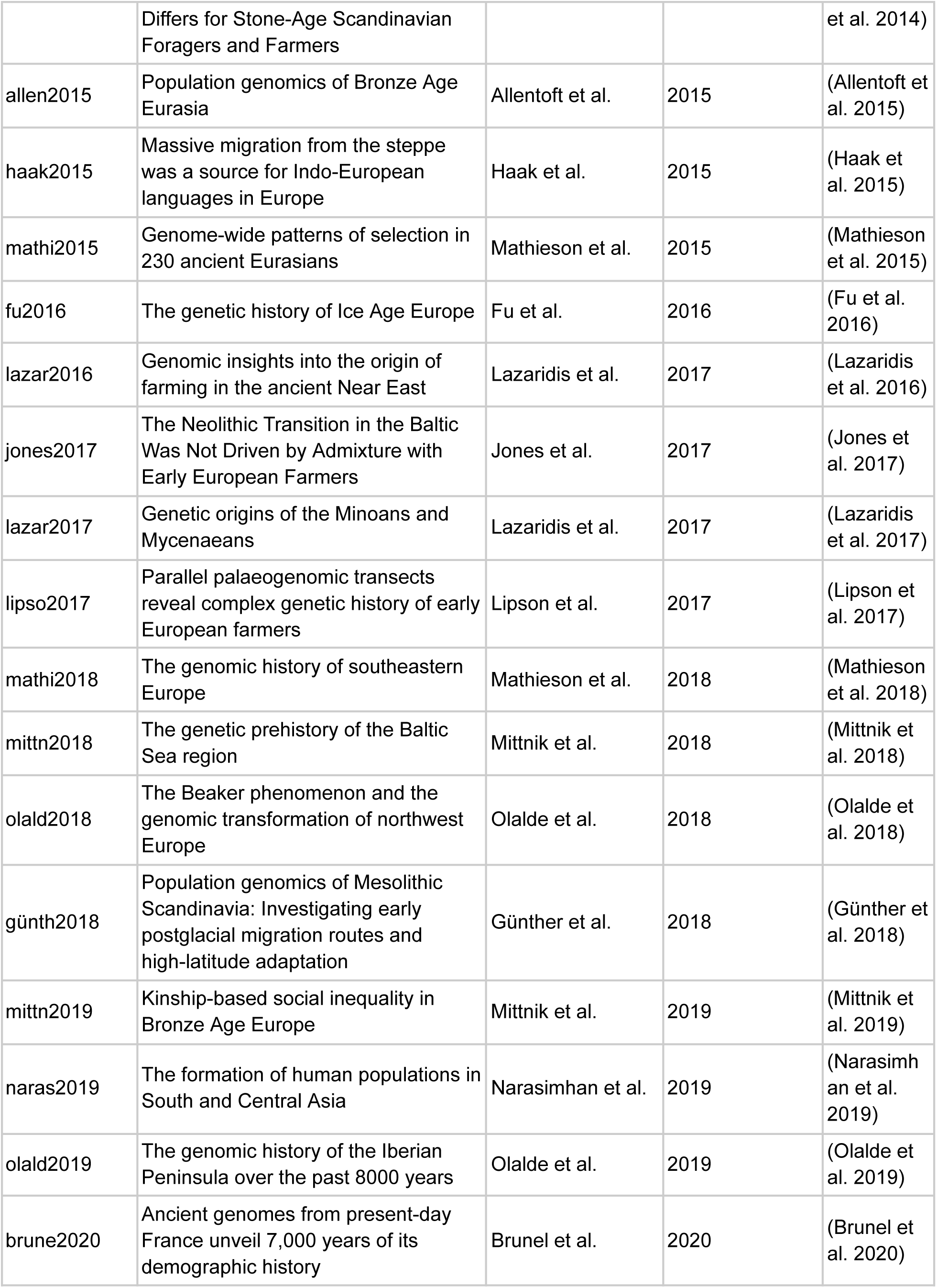
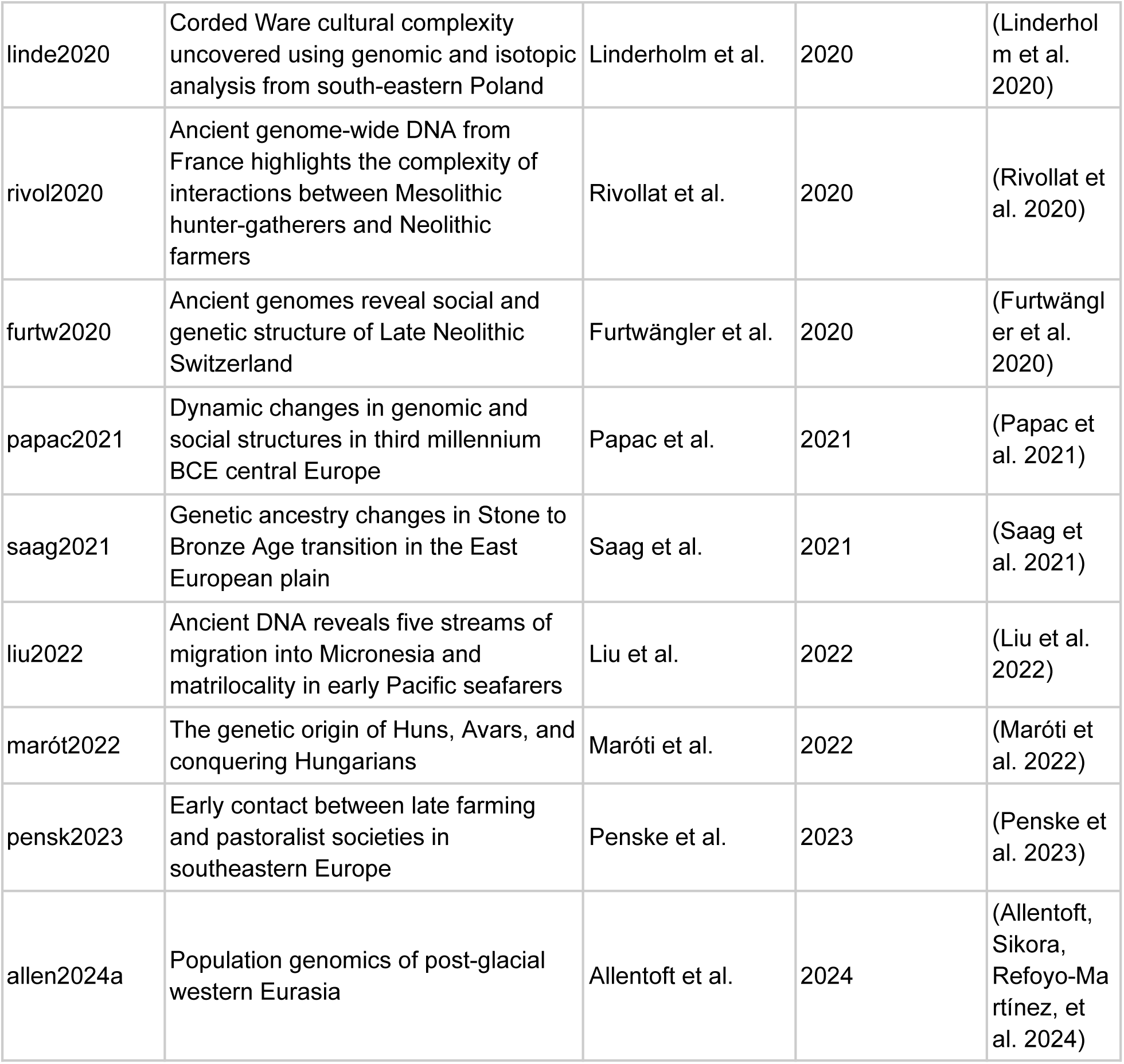
Human paleogenomic studies sampled in this study.

**Table 2.** Archaeological metadata and candidate explanatory variables used in the study.

| Data type | Tier | Variable | Description |
| --- | --- | --- | --- |
| Archaeological metadata | Core | Geolocation | Precise site location reported as coordinates |
|  |  | Individual ID | Unique identifier of the archaeological individual in the source publication |
|  |  | Multiple names | Other names the site was reported as, including local languages and alphabets |
|  |  | Overall chronology | The entire documented chronological sequence of the site |
|  |  | References | References to previous publications of the site, findings, or analyses |
|  |  | Sample chronology | Dating of the sampled individual |
|  |  | Site description | General summary of findings from the excavated area |
|  | Extended | Anthropological age | Biological age based on physical anthropology; synonymous with age-of-death |
|  |  | Anthropological sex | Biological sex based on physical anthropology |
|  |  | Body arrangement | Arrangement of the individual according to the cardinal directions |
|  |  | Body positioning | How the body was placed inside the grave |
|  |  | C14 | Reported radiocarbon dating |
|  |  | Culture | Cultural affiliation of the sampled individual |
|  |  | Grave | Information about the immediate find circumstances of the individual; synonymous with context |
|  |  | Grave goods | Finds discovered with the individual |
|  |  | Images | Images showing the grave, individual, and/or grave goods; synonymous with accompanying finds |
| Explanatory variable |  | Archaeology Index | Average disciplinary affiliation profile of all authors |
|  |  | Number of authors | Total number of authors |
|  |  | Number of countries | Total number of countries from which the samples were reported |
|  |  | Number samples | Total number of newly published and re-sequenced sampled individuals |
|  |  | Year of publication | Year the study published in the journal |

We find that archaeological metadata in these studies has been reported haphazardly. We conclude that, over the last 11 years, the most conspicuous improvement has occurred in the reporting of a sample’s Geolocation but that variation in reporting for the other variables is generally not well explained by the factors we tested. In light of our findings, we propose a metadata reporting checklist to help researchers structure the reporting of archaeological metadata. This structure could potentially constitute the basis for a future, more robust solution.

## Results

Our dataset comprises 30 publications, covering 3911 newly published or re-sequenced ancient human individuals from 49 countries with a predominant focus on Western Eurasia (Fig. 1). The number of samples per publication varied from 2 to 522 with a mean of 130.37.

**Figure 1:**
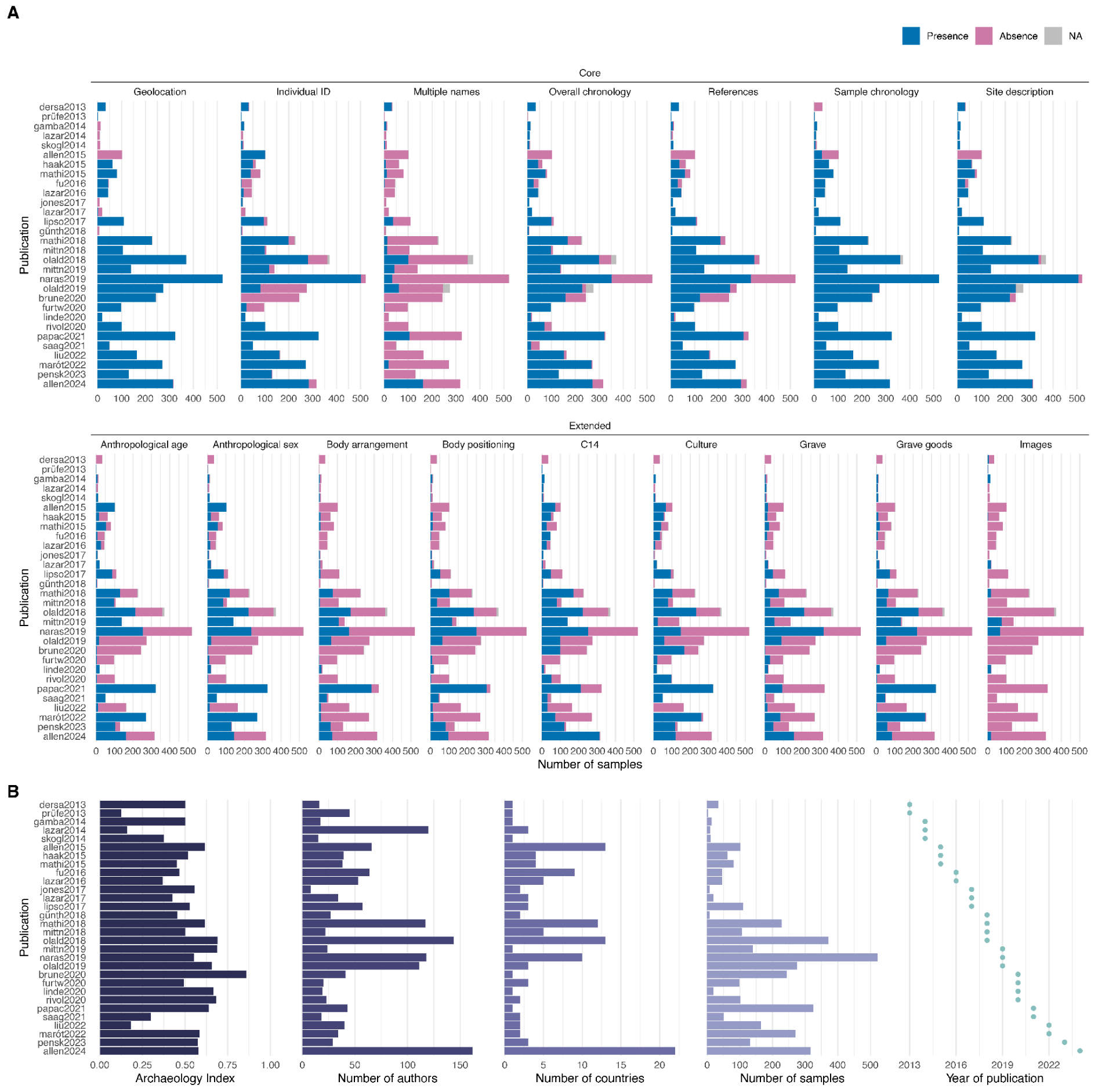
Metadata and candidate explanatory variables. (A) An overview of archaeological metadata reporting for all sampled individuals of the 30 peer-reviewed publications from 2013 to 2024. (B) An overview of the candidate explanatory variables

For each sample, we recorded the availability of 16 metadata types typically expected in an archaeological publication (Table 2). The variables selected for the study derive from an on-going collection of archaeological data for the COREX project (https://www.gu.se/en/research/corexfrom-correlations-to-explanations), specifically the cultural data associated with individuals sampled for aDNA research. The variables selected for the purpose of the project correspond to the generally accepted standards of publishing archaeological individuals in the scientific literature (Parker Pearson 1999; Nilsson Stutz 2003; Nilsson Stutz and Tarlow 2013; Tõrv 2016; Knüsel 2022; Reiter et al. 2024) and are routinely used for relative dating, cultural classification, or investigation of intercultural ties between contemporary or consecutive communities. It should be stressed that some degree of discrepancy caused by the evolution of the discipline, divergent archaeological schools, discovery circumstances and preservation of finds, and last but not least, culture-specific depositional behavior, is to be expected. However, it is because of the standardized data protocol we have developed that problems with reporting archaeological data were encountered, and after extensive documentation over a period of 2 years, the initial protocol was re-worked to prepare a template for the evaluation of archaeological metadata reporting standards in paleogenomics.

The data collection protocol was based on a hierarchical structure where the highest order is the archaeological site and archaeological information recorded for it which nests the sampled individual in time and space. Other variables were specifically related to the sampled individual, like discovery circumstances or cultural traits specific for archaeological research. By including both levels of information it is first possible to discern between individuals found in different locations, and then between individuals recovered in the same location.

These variables fall into either ‘core’ or ‘extended’ tiers based on their functions. The core tier comprises metadata that serve basic identification purposes; they describe the “who”, “when” and “where” of a sample. The extended tier provides further information on bioanthropological determinations and cultural traits specific to the sampled individual.

For each of the 3911 sequenced humans, we recorded the availability of information for each metadata variable. Note that availability here refers to the presence or absence of reporting rather than of the data itself (e.g. reporting of missing data is considered as present). In cases where presence could not be determined from the publication, “not applicable” (NA) was assigned (see <u>Materials and Methods</u>). These represent only 0.52% of observations overall. We present this archaeological metadata reporting dataset in Figure 1, illustrating the presence/absence status for each sample published in the examined studies.

To quantify variation in reporting, we aggregated the presence/absence observations per sample, i.e. 1 (presence), 0 (absence) or 0.5 (NA), into a percentage presence (completeness) for each publication, i.e. [0,1], for each metadata variable (Fig. S1). This allowed us to treat every publication with equal statistical weight, and therefore retain an effective sample size of 30. For clarity, we denote the variables in bold and italic.

We found significant differences in the level of completeness across metadata variables (Kruskal-Wallis test χ^2^(15, 480) = 199.76; p < 2.2e−16). Core variables tended to have greater completeness than the extended variables (post-hoc Dunn tests Fig. 2A; Table S6), with *Site description* being the most complete. Our dataset also showed that core variables generally had a lower completeness variance than the extended variables, except for *Geolocation* and *Individual ID*, which had relatively high variances among all metadata (Fig. 2A). In addition, we found that the variation in completeness is more influenced by differences in the type of metadata (Kruskal-Wallis η^2^ = 0.398, 95% CI [0.34-0.50]) than by the differences across publications (Kruskal-Wallis η^2^ = 0.080, 95% CI [0.07-0.21]).

**Figure 2:**
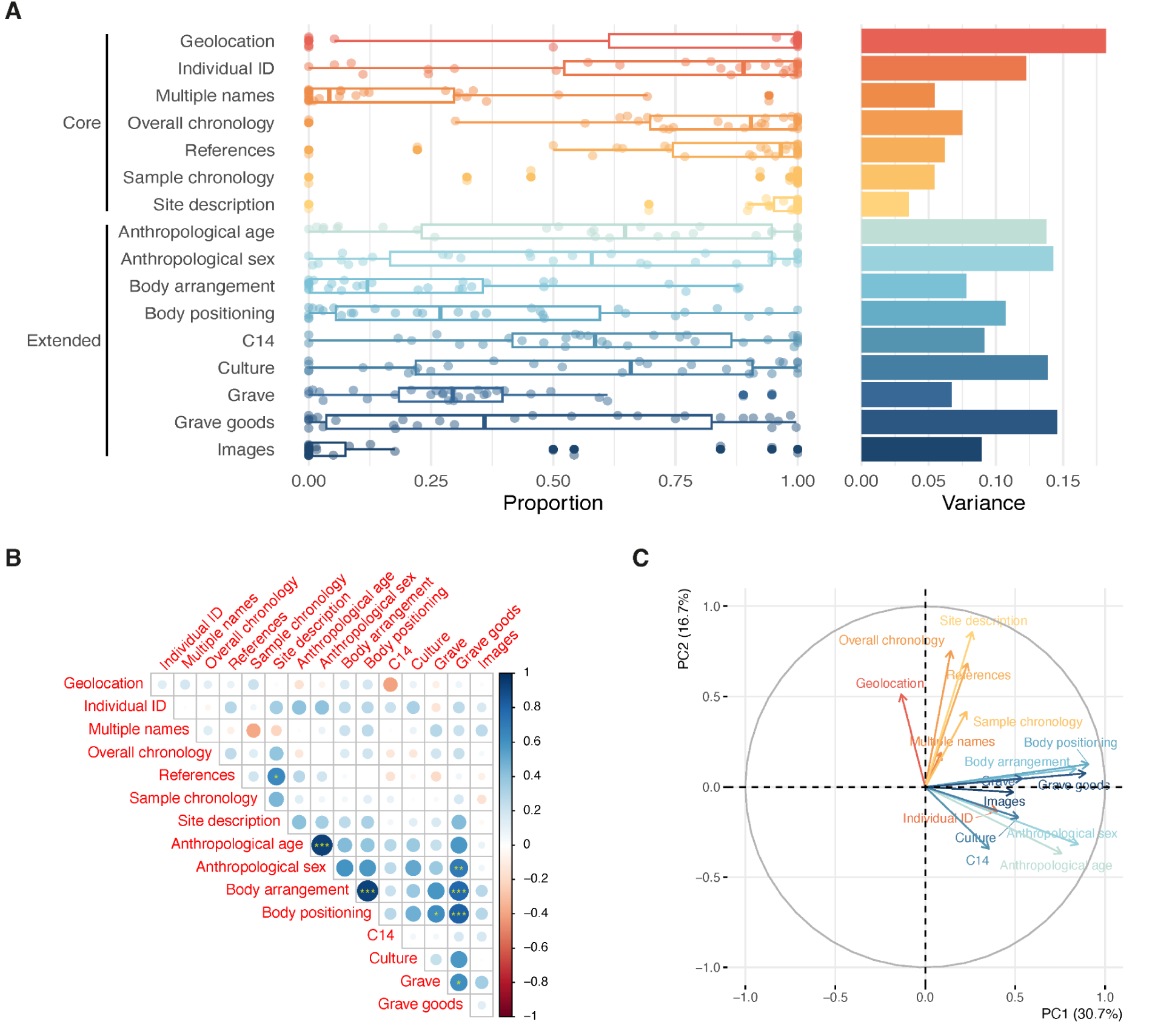
Variation and correlation in metadata completeness. (A) The box plot shows the distribution of the data which are represented by the points. Each point corresponds to the proportion of a metadata variable for a given publication. The bar plot illustrates the sample variance of the proportions. (B) Spearman correlations. Spearman’s ρ is indicated by the colour and the size of the circle whereas the significance level after Bonferroni correction is indicated by the asterisks (“*”: padj < 0.05; “**”: padj < 0.01; “***”: padj < 0.001). (C) The loading plot of PCA. The projected vectors correspond to the contributions of metadata variables to the PCs as measured by correlations.

Next, we analyzed correlations between metadata variables, in their level of completeness. We first calculated the Spearman’s rank correlation (ρ) between each pair of the 16 metadata variables. Generally, correlations were weak (non-significant accounting for multiple testing). However, we found significant positive correlations between demographic and burial-related categories respectively (Fig. 2B).

To better understand the overall variance, we performed a principal component analysis (PCA) to identify if there was any correlation structure in the completeness of variables (Fig. 2C). Briefly, a *m × n* matrix of completeness was transformed into a new coordinate system of *m × p*, where *m* denotes the publications, *n* the metadata fields and *p* the principal components. Specifically, we were interested in how different metadata contributed to the principal components. We found that PC1 and PC2 explained 30.7% and 16.7% of the total variation, respectively, and the clustering in the loading vectors supported and more clearly represented the pairwise correlations previously found. Additionally, we found that the first two PCs broadly correspond to the extended and core metadata categories, respectively (Fig. S8G-H).

To identify whether some basic structural factors would influence the completeness of reporting, we considered five plausible explanatory variables: (1) the proportion of authors with a primarily archaeological background; (2) the total number of authors; (3) the number of sequenced samples; (4) the number of countries from which the samples originate; and (5) the year of publication (Fig. 3). They were chosen because they are quantifiable and comparable variables that capture the scope and bibliographic information of the studies. As an estimate for (1), we devised an affiliation-based metric we call the “Archaeology Index” that aims to approximate the contribution of archaeologically-trained researchers in a given study (see <u>Materials and</u> <u>Methods</u>). We recognize that representing author backgrounds as a binary categorical variable (archaeology, not archaeology) does not capture the full range of researchers’ expertise, but as a first order of approximation, institutional affiliation should give a good proxy for expertise. In addition, we log-transformed the observations of candidate explanatory variables (2)-(4) as they appeared to be skewed (Fig. S4).

**Figure 3:**
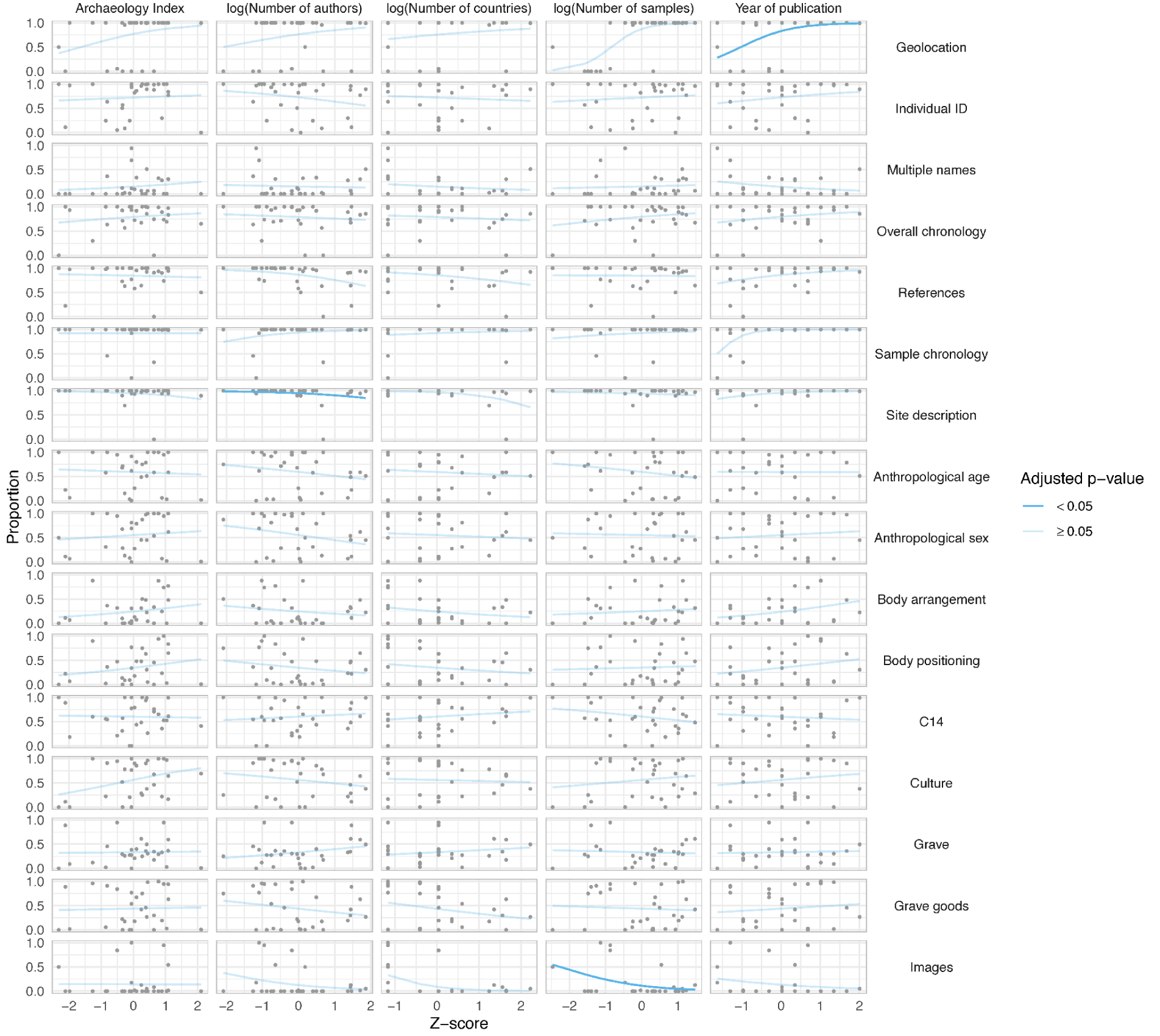
Model fit of fractional logistic regression model. An opaque line indicates the coefficient estimate (β^) is significant after Bonferroni correction.

To determine whether these candidate explanatory variables can explain variation in the completeness of the metadata response variables, we used fractional logistic regression, which is suited to working with bounded fractional response variables (see <u>Materials and Methods</u>). Generally, the explanatory variables provided very little explanatory power (Fig. 4A). Across all models, we only identified a statistically significant positive effect for the year of publication on *Geolocation* after adjusting for multiple testing (β^ = 1.56, padj = 0.018), suggesting improved reporting of geographical coordinates over time. We also identified negative associations between Images and the log(number of samples) (β^ = −0.92, padj = 0.017) and between *Site description* and the log(number of authors) (β^ = −0.65, padj = 0.024), suggesting these two metadata variables have lower completeness in the larger publications that sequenced more ancient humans and had a longer author list.

**Figure 4:**
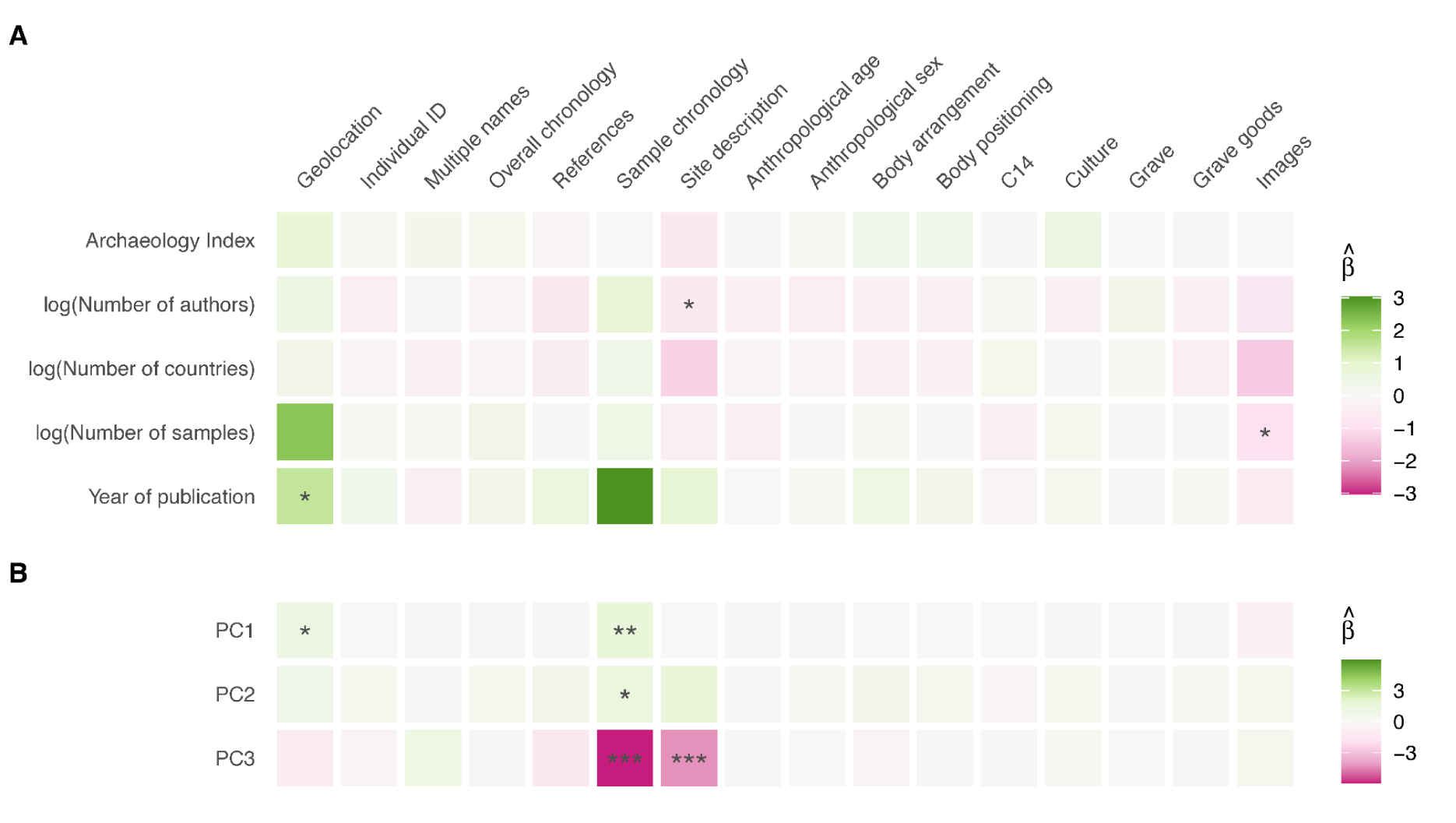
Model results. (A) The effects of candidate explanatory variables on metadata completeness. The coefficient estimate (β^) is indicated by the color, whereas the significance level is indicated by the asterisks (“*”: padj < 0.05; “**”: padj < 0.01; “***”: padj < 0.001). (B) The effects of candidate explanatory variables on metadata completeness. The coefficient estimate (β^) is indicated by the color, whereas the significance level is indicated by the asterisks (“*”: padj < 0.05; “**”: padj < 0.01; “***”: p_adj_ < 0.001). Only the significance levels from the models significant against the null model based on the Wald test are shown.

In an attempt to identify latent explanatory variables – not well represented by our individual variables, but perhaps captured by a combination of them – we carried out a PCA of the five explanatory variables, and then performed a multiple regression analysis using the first three PCs (explaining 90% of the variation in all five explanatory variables (see <u>Materials and</u> <u>Methods</u>). Across all metadata, only *Geolocation* (χ^2^(3, 26) = 8.51, p = 0.037), *Sample chronology* (χ^2^(3, 26) = 14.12, p = 0.003), and *Site description* (χ^2^(3, 26) = 13.21, p = 0.004) were statistically significant based on the Wald test. In these models, PC1 appears to have significant positive associations with *Geolocation* (β^ = 1.19, padj = 0.019) and *Sample chronology* (β^ = 1.51, padj = 0.002); PC2 has a positive association with *Sample chronology* (β^ = 1.37, padj = 0.01), while PC3 has a negative association with *Site description* (β^ = −4.23, padj = 0.001) (Fig. 4B). We note, however, that here PCs are linear combinations of the explanatory variables, and that the regression is simply a test for (linear) association (Fig. S11). Overall, *Geolocation* and *Site description* were the only two metadata variables found to be impacted by at least one of the explanatory variables in both simple and multiple regression analyses.

## Discussion

Our pilot study found considerable variation in the completeness of metadata reporting across variables and publications. We identified improvement in the completeness of *Geolocation* over time, but our candidate explanatory variables poorly explained differences in other variables. Rather, it is likely that the lack of a universally accepted standard for reporting cultural data in archaeology is a significant contributing factor to this (Reiter et al. 2024). In the absence of clear guidelines, archaeologists make individual decisions shaped by country of origin, period, school, and experience, which collectively generate an impression of inconsistent data reporting. It might also be influenced by the internal dynamics of the research group, such as the mode of collaboration between researchers from different disciplines and their differential contributions to the final publication, which are not captured by the broader quantitative variables we chose.

The informal conversations we carried out with researchers working in both small and large interdisciplinary teams have often revolved around tensions emerging during sampling and writing. The archaeologists we spoke to often expressed the view that the guidelines for supplementary information on the sampled individuals were vague and imprecise. Simultaneously, the ambiguous requirements were on occasion accompanied by explicit anticipations of straightforward classifications, e.g. “Is this individual Linearbandkeramik?” or “Is it Corded Ware?”

Paleogenecists tended to respond to this issue with discomfort as, in their view, they considered archaeologists they worked with experts in their field, better suited to make informed decisions on what the relevant level of detail is. At the same time, paleogeneticists frequently encountered problems with requesting their collaborators to provide any supplementary information at the writing stage. This criticism tended to be relegated back to the unclear guidelines the archaeologists received from their collaborators.

Beyond authors’ anecdotal reflections, collaboration dynamics between archaeologists and geneticists have been a subject of discussion concerning the interdisciplinarity of the field. A common theme that arises is the asymmetric relationships that privilege genetic evidence over that of the humanities disciplines due to various conceptual, epistemological and methodological gaps, as well as structural issues (e.g. funding and publication incentives) that promote them (Hofmann 2015; Horsburgh 2015; Vander Linden 2016; Heyd 2017; Ion 2017; Callaway 2018; Furholt 2018; Booth 2019; Charlton et al. 2019; Frieman and Hofmann 2019; Sykes et al. 2019; Downes 2021; Källén et al. 2024; Mulcare and Pruvost 2024). They have long been raised in the contexts of the sampling process, research questions, data analysis and interpretations. However, without a comprehensive survey-based evaluation (Källén et al. 2024), it remains unclear how they contribute to metadata reporting despite journals’ requirements to publicly disclose any conflicts of interest and individual contributions. Furthermore, while these factors affect what kind of data goes into a publication, we argue metadata reporting is a broader issue in itself as part of the data management and stewardship discussion (Wilkinson et al. 2016), as exemplified by the data archiving developments in genomics (Field et al. 2008; Yilmaz et al. 2011; Medema et al. 2015; Bowers et al. 2017; Roux et al. 2019; Bergström 2024; Karsch-Mizrachi et al. 2025). Nonetheless, it seems that this aspect has been overlooked in the critiques and best-practice guidelines to date (Sirak and Sedig 2019; Wagner et al. 2020; Alpaslan-Roodenberg et al. 2021), even though archaeological data – whether they are directly used in the analysis – describe the individuals and lend support to archaeological interpretations.

The observed improvement over time in *Geolocation* could be due to the increasing need for spatiotemporal information to relate genetic processes to other processes in space and time (Loog et al. 2017; Racimo, Woodbridge, et al. 2020; Schmid and Schiffels 2023). As a result, these metadata became increasingly visible in the SI, even though they have always been mandatory for submission to genomic databases such as the European Nucleotide Archive (ENA), together with sample age and sample type/material (e.g. petrous bone and tooth) (Charlton et al. 2019; Bergström 2024; Bergström et al. 2024). Nevertheless, these databases have not extended their schema to incorporate the rapidly growing aDNA sequencing data. This led to the creation of aDNA data repositories such as the Allen Ancient DNA Resource (AADR) (Mallick et al. 2024) and Poseidon (Schmid et al. 2024) and sparked the emergence of aDNA communities such as the Standards, Precautions, and Advances in Ancient Metagenomics (SPAAM) community (https://www.spaam-community.org/) and the Minimum Information about any Ancient Sequence (MInAS) project (https://www.mixs-minas.org/) which spearhead the development of community standards in response to the challenges in aDNA data sharing and archiving (Fellows Yates et al. 2021; Bergström et al. 2024). We believe these factors have likely resulted in the widely established aDNA metadata fields that report the most basic information about the samples.

We identified a decrease in the completeness of *Images* reported as the number of samples increased, which likely represents the difficulties in acquiring permits for reproducing existing images of burials and finds in papers with a large number of new sequences. As paleogenomic studies generally use samples from already reported discoveries, this problem is usually resolved by providing references to original publications. We found a relatively high completeness level for *References*, suggesting that this information may be obtainable from the source publications. However, the process of acquiring and examining each publication can be challenging, time-consuming, and the source publications may not contain relevant images. Since images are as close as possible to raw archaeological data, indicating whether they can be obtained in published resources can streamline data collection for other researchers.

We also identified a decrease in *Site description* reporting as the number of authors increased. This information allows correct attribution of samples to their origins and facilitates differentiating between locations with similar names that are found in immediate proximity. A plausible explanation is that it becomes more challenging to assemble all the necessary metadata in the SI as the scope of papers increases and requires more extensive collaborative work. As the length of supplementaries increases to accommodate hundreds of sites with single sampled individuals, the likelihood of erroneous omission of a description increases. It is possible that these errors are also caused by a poor design of the compilation workflow in which a single researcher is responsible for ensuring that hundreds of descriptions find their way into the SI. Given that a sample list has to be also prepared for the tabulated SI, implementation of a standardized *Site description* would ensure that such errors are mitigated. Currently, the peer-review process encourages but does not require the evaluation of supplementaries, which raises the question whether submissions undergo sufficient quality control.

Apart from these widely established aDNA metadata fields, *Sample chronology*, *Overall chronology*, and *References*, show consistently high completeness across publications. This trend only partially overlaps with the metadata fields of AADR (Mallick et al. 2024), Poseidon (Schmid et al. 2024), or recommendations from the different metadata consortia. The relatively high variance of Individual ID reporting could result from assigning sample IDs to aDNA data, deeming Individual ID from the original archaeological literature optional. *Multiple names*, which records inclusion of original site names from countries using non-Latin alphabets, was consistently missing from the core metadata. This could pose severe challenges for tracing originally published contexts.

Unlike core metadata, the extended tier is not well scrutinized at the peer-review stage, and efforts to collect large archaeological datasets remain specific to particular research projects (Reiter et al. 2024). At the time of writing, large databases that incorporate archaeological data from different studies or metadata consortia addressing the data reporting standards are in their infancy. This, in turn, makes it challenging to openly propose which data carries the most importance and which form of reporting is most effective. However, the increasing rate at which archaeological data is collected and reported in paleogenomic papers raises the question of whether it is time for these discussions to emerge to counteract the reporting variability we document here. Its importance cannot be overstated as the increasing number of published paleogenomes is already causing problems with correct attribution of samples to particular individuals, distinguishing between sequenced genomes originating from the same site, and distinguishing duplicates from twins. This is an avenue where archaeological data will prove important, as the combination of variables should reduce misattribution or duplication of results. The currently utilized standard that relies on geolocation, sample date, and sample type is insufficient as often more than a single sequenced genome originates from the same archaeological site, same grave, or even the same individual.

Beyond challenges with data attribution, the higher variability in metadata reporting has broader impacts on the interdisciplinary nature of human paleogenomic and archaeological research. Without explicit reporting standards, exploratory or explanatory research on links between patterns of genetic and cultural changes will not only have to face conceptual or theoretical challenges but also be hindered by data collection required for empirical analysis. The time-consuming and error-prone process of individual tracing of every relevant publication to obtain information that is not necessarily the most up-to-date can be streamlined if clear guidelines are provided to data curators responsible for providing samples for paleogenomic research. As archaeologists and curators with access to the most detailed descriptions of archaeological finds, either through excavating the individuals themselves or working in museums (Horsburgh 2015), they are the most competent to report the fine-grained details of the samples they have acquired. If provided with a clear outline of what should be considered the minimum required information, they can create standardized descriptions to be reevaluated in the future. Lastly, it would address the ever-persistent challenge of heterogeneous archaeological data reporting standards allowing archaeologists to more effectively utilize the data reported in aDNA papers in their work (Alpaslan-Roodenberg et al. 2021; Reiter et al. 2024). This is crucial for a more detailed exploration of the relationship between genetic and cultural patterns. Booth et al. recently highlighted the relevance of high-quality reporting standards by critically examining SI to provide a broader context for the genetic turnover patterns in the British Isles during the Bell Beaker period (Booth et al. 2021).

A change in the existing metadata reporting standards has to be adopted by all stakeholders. Although we cannot rule out potential bias towards several large institutions in the field and supraregional practices given our sampling scope for macroscale studies on Western Eurasia, aDNA sampled individuals that fall within the scope are overrepresented in the global dataset (Mallick et al. 2024) and hence deserve considerable attention. As paleogeneticists have already created consortia directed towards developing a better metadata standard for aDNA sequences, initiatives similar to MInAS directed towards the structuring of archaeological metadata are needed (Reiter et al. 2024). Our results indicate that although authors affiliated with archaeological institutions were always part of the research groups and discussions throughout the paleogenomics revolution, insufficient attention was paid to the archaeological data ending up in the final publications and SI. Here, we echo with Horsburgh (Horsburgh 2024; see reply to Källén et al. 2024) that archaeologists have the leverage and power to engage more meaningfully in human paleogenomic studies and metadata reporting should not be overlooked, alongside all the ethical and interdisciplinary discussions. Otherwise, generations of researchers will find themselves struggling with an incomplete and arduous dataset to conduct novel investigations into the human past.

The data availability issues we have identified can be structurally overcome in the future through the implementation of mandatory data import to existing or currently-in-development repositories or databases designed according to FAIR and CARE principles. Such infrastructure will have to tread carefully between requiring detailed information about the archaeological individual while allowing researchers to specify data unavailability or restrictions about disclosing confidential information where necessary. Presently, determining which of the two cases applies to the missing data we have documented can only be applied retrospectively and at great cost in labor and time. However, given the accelerating rate of publishing paleogenomic research, an intermediate and easy-to-implement solution is the necessary first step to improve the on-going reporting and provide sufficient information to estimate which metadata fields should eventually become optional.

## Checklist for reporting archaeological data

Drawing on the work of communities involved in evaluating and improving standards in publishing aDNA data (SPAAM and MInAS), from the results of our analysis, and from conversations with researchers, we propose a minimum information standard for reporting archaeological data in human paleogenomic research. We acknowledge that the archaeological record is complex, shaped by the history of the finding as well as skills, knowledge, and access to technology archaeologists had at the time of discovery. A full-fledged, comprehensive reporting standard will require a structured digital solution with explicit and accessible vocabularies developed through discipline-specific consultations and discussions of stakeholders in human paleogenomic research. We believe our checklist is only a first step towards realizing this goal, firstly, by indicating where gaps already exist; secondly, by proposing a tentative list of fields that are routinely provided in archaeological literature; thirdly, by indicating where common grounds for integrative reporting already exist.

To construct a systematic reporting standard, unique identifiers are necessary to link data generated from different sources. When it comes to human paleogenomics, the majority, if not all, of the studies have been based on aDNA extracted from archaeological individuals. In these cases, an individual is the unique identifier and the natural bridge between aDNA and archaeological data. Therefore, we believe that the archaeological individual – the specific, archaeologically discovered human being – who was found, documented, and reported in scientific research should be adopted as the primary data reporting unit. This standard of data reporting allows all the relevant archaeological and genetic data to be linked to a single, unique individual, providing a more robust way of associating different strands of relevant data. It offers the advantage of preventing data duplication or misattribution, which is especially important as the number of available genomes continues to increase rapidly, requiring more time and effort investment into aggregation and harmonization of datasets (Wilkinson et al. 2016; Mallick et al. 2024; Karsch-Mizrachi et al. 2025). This standard was adopted by the COREX project and its Big Interdisciplinary Archaeological Database (BIAD) (Reiter et al. 2024) to enable a more streamlined data collection protocol and limit the possibility of data duplication or misattribution.

We believe other researchers will benefit from it and can improve upon it by an informed consideration of how to best report specific cultural data. With the right kind of momentum and increasing data availability, we hope that initiatives like SPAAM or MInAS, will emerge around this data standard and provide discussions to further improve data reporting. The responsibility for collecting cultural data falls on the archaeological community, as archaeologists are the experts in their field. However, ensuring that this information is provided in a comprehensive form will be of benefit to archaeologists and geneticists alike.

Presently, following our experience with associating paleogenomic data with archaeological individuals, we propose the following checklist for reporting archaeological data from aDNA-sampled individuals, and crucially, we recommend this to be checked at peer review (Table 3). An example of how such a tabulated metadata reporting style could look is included in the SI (Table S7).

**Table 3:** Checklist of archaeological metadata.

| Level | Data type | Description | Details to include |
| --- | --- | --- | --- |
| Site | Name | Site name | Historical names |
|  |  |  | Alternative reporting conventions |
|  |  |  | Local scripts, transliterations, and diacritic signs |
|  |  |  | Heritage board identification number |
|  | Geolocation | Site location | Degrees decimal |
|  |  |  | Report measurement uncertainties |
|  |  |  | If cannot be disclosed for heritage protection reasons, state that information explicitly |
|  | Type | Site type | Functional identification of the discoveries |
|  |  |  | If the site is phased and multi-cultural provide information about the changes of function through time |
|  | Dating | Chronological characterisation of the site | Specify dating information, including how it was generated |
|  |  |  | If the dating was established using radiocarbon dating, include lab codes, material type, and raw uncalibrated data |
|  |  |  | If archaeological dating was provided, include the method and criteria used for its estimation |
|  |  |  | If possible, include quantification of the documented findings |
|  | Culture | Cultural classification of the site | If archaeological cultures are not reported in the area, make that information explicit |
| Individual | Name | Individual's name as reported in the first publication | Include alternative reporting conventions |
|  |  |  | Include the lab ID used for genetic sequencing |
|  | Dating | Chronological positioning of the individual | If absolute dating was obtained using archaeometry, include lab codes, material type, and raw (uncalibrated) data |
|  |  |  | If archaeological dating was used, report the method and criteria used |
|  | Find circumstances | Context of deposition | If the individual comes from a grave, report details of the grave as well such as type, shape, and dimensions |
|  |  |  | Specify if the individual was discovered in a collective grave |
|  |  |  | If the individual comes from a different context (settlement, random discovery) provide it with as much details as possible |
|  | Culture | Cultural classification of the individual | Report the cultural group and the classification method |
|  |  |  | If the individual originates from a region or is assigned a time period without cultural groups, be explicit about this |
|  | Burial rite | Burial rite the individual's body was subject to | Inhumation |
|  |  |  | Cremation |
|  |  |  | Partial cremation |
|  | Body position | Position of the body in its context | Crouched |
|  |  |  | Extended |
|  |  |  | Supine |
|  |  |  | Sitting |
|  |  |  | If indeterminate, state this explicitly |
|  |  |  | If possible include the image of the individual *in situ* |
|  | Body orientation | Orientation of the body in relation to world directions | Angles |
|  |  |  | Cardinal directions |
|  |  |  | If indeterminate, state this explicitly |
|  | Anatomical completeness | Body treatment of the deceased | State whether the individual's body was manipulated post-mortem |
|  |  |  | Specify if such an evaluation was not performed or other factors prevented such an analysis |
|  | Anthropological sex | The osteological sex of the sampled individual | If the analyses yielded an uncertain classification or proved indeterminate, include this information |
|  |  |  | Uncertainty can be reported as confidence probability, where 0 represents 100% confidence of male determination, while 1 represents 100% confidence of female determination |
|  | Anthropological age | The osteological age of the sampled individual | If the analyses yielded an uncertain classification or proved indeterminate, include this information |
|  |  |  | If no evaluation took place include this information as well |
|  | Accompanying finds | Artefacts discovered alongside the sampled individual | If possible, provide images |
|  |  |  | If finds have been classified following an existing typological scheme include the reference |
|  |  |  | If the finds have been published in detail elsewhere provide a reference |
|  |  |  | If no accompanying finds were documented, state that explicitly |
|  | Health markers | Bioanthropological evidence of the ante-mortem health condition of the individual | Include an explicit statement about absence of documented health markers |
|  |  |  | Report if such information is being prepared for a separate publication |
|  | Images | Raw data allowing re-classification of findings | Find circumstances |
|  |  |  | Body position |
|  |  |  | Body orientation |
|  |  |  | Accompanying finds |
|  |  |  | Health markers |

We encourage authors to include images or hyperlinks to images with find circumstances, body positioning, body orientation, and grave goods, as (1) they allow visual identification of the sampled individuals to rule out misattribution or duplication of aDNA sequencing; (2) they are the closest to “raw” data on cultural variables associated with the sampled individuals; (3) they provide the possibility of verifying and reclassifying data. In the past, the publication of archaeologically-documented human remains and accompanying finds preceded interdisciplinary analyses, providing a strong basis for the identification of archaeological individuals. As paleogenomic research becomes more widespread and valued due to prestigious and high-impact research papers, there is a growing number of previously unpublished archaeological individuals which have little if any archaeological data associated with them. These individuals play a significant role due to the association with aDNA sequences but have limited archaeological relevance due to the missing information.

Simultaneously, we acknowledge the objective difficulties in acquiring permissions for republishing images from existing sources, especially for previously published individuals, and recommend that, in these instances, a source reference should be provided in the SI. When it comes to issues with supplementary size limitation, one potential solution is to apply a unified naming protocol to the digital images (file names and text labels) and to upload them to a public repository like Zenodo (European Organization For Nuclear Research and OpenAIRE 2013), referencing the publication. We recognize that in some countries the presentation or publication of images of the dead is banned or prevented by publishing outlets. Finally, we realize the ethical challenges involved in reporting archaeological data and remaining sensitive to laws and customs of aboriginal people. In cases where providing detailed cultural data would infringe on these rights, an explicit statement would make restricted data sharing more transparent. We encourage authors to explicitly state these conditions in the SI, so that the researchers attempting to incorporate such data in their work can account for the cultural preferences and legal regulations. In all instances absence of information about any of the aforementioned variables should be explicitly stated to prevent subjective data input. While such a standard has been already encountered in one of the analysed papers (Papac et al. 2021), it should become the default for reporting archaeological data.

We are confident that the checklist will serve as a template for reporting archaeological data that will streamline the publication of ancient DNA research using prehistoric and historic individuals in the future. In the long term, we envision a community-driven checklist providing an explicit set of requirements that paleogeneticists should expect from their collaborators. Secondly, irrespective of their temporal or regional interests, archaeologists will be able to assess which information should be included in such a report. Thirdly, reviewers will know which information they should seek in the SI files and what to mark as incomplete or missing entries.

## Conclusions

The paleogenomics revolution in archaeology is here to stay; whether it will spearhead truly interdisciplinary research on the human past will depend on our ability to effectively integrate rich archaeological data with genetic data. Collaboration and communication are the necessary first steps. Adopting strict practices of academic rigor in data reporting and curation can be challenging, but it will enable more complex research questions to be addressed, further blurring the line between the humanities and sciences (Racimo, Sikora, et al. 2020).

As cultural information has been reported in an unstructured form since 2013, our recommendations do not require creating a new workflow as much as modifying the already existing one and allowing geneticists, archaeologists, and reviewers to address the quality of information they use in their research. This is why the key structural component is to ensure that the sample metadata reporting undergoes evaluation as part of the default manuscript review process. We are confident that, in our examination of the first 15 years of paleogenomics, we have identified a critical issue in an otherwise cutting-edge research field. In order to make the paleogenomics revolution in archaeology truly revolutionary, this issue needs to be tackled ahead of time for the benefit of all parties involved.

## Materials and Methods

### Sampling of papers for metadata collection

We sampled 30 peer-reviewed articles (Table 1) published between 2013 and 2024, as this period represents the most intensive period of aDNA analysis in archaeology. No regional or chronological preference was given, although the prevalence of studies on Eurasian prehistory is observable, as it is the area and period most frequently chosen in evolutionary genetics (Roca-Rada et al. 2020; Mallick et al. 2024; Sawchuk et al. 2024; de la Fuente Castro and Figueiro 2025). However, given the high density of sampling of this space and time it should be considered representative for determining the quality of the reported archaeological data and estimating the overall reporting standards. We sampled at least 2 papers per year published since 2013 and initially included pre-prints. We focused on macroscale studies with a large number of individuals because, unlike site-specific studies in which archaeological data are often integrated and examined in the main text, they usually report the data as metadata in the SI. The data collection was finalized in early 2023 with a single paper (Penske et al. 2023). While preparing the paper one of the pre-prints was published (Allentoft, Sikora, Refoyo-Martínez, et al. 2024; Allentoft et al. 2022 Jan 1), so the paper was re-examined to reflect the peer-reviewed version and bibliographic reference was modified to reflect the final version of the study.

### Data collection

The descriptive SI was the primary source of data since it is the designated section for reporting details about the sampled individuals. The tabulated SI files were examined to extract information about the coordinates, radiocarbon dating, or sexing of the individuals. However, cross-referencing the two sources has often revealed the discrepancies between the genetic identification numbers reported between SI of the same paper. Most commonly encountered problems were aggregations of IDs reported in two or more columns, reordering of the ID numbers, or replacement of symbols, such as hyphens or backslashes. For convenience, it was decided to rely on the ID numbers reported in the descriptive SI as they correspond directly to the archaeological descriptions. The varying standards and qualities of reporting archaeological data in the descriptive SI meant that the data collection had to be handled manually.

### Data preparation

The reporting of archaeological metadata was classified as “presence”, “absence” or “not applicable” which were encoded numerically as 1, 0, and 0.5. The latter applied to metadata of previously published individuals re-sequenced in the publications examined or those of new individuals from sites that were reported in previously published studies. As only the publications listed above were examined, we considered 0.5 fair since we had no information on the presence or absence of the metadata. We also provide a more nuanced classification of the data in the Supplementary, which for the final analysis was reduced to presence or absence, irrespective of the overall quality. This data was transformed into proportions per publication for downstream quantitative analyzes.

### Archaeology Index

The Archaeology Index was used to measure the contribution of archaeology-affiliated authors. To minimize the subjectivity of assigning author affiliation due to limited access to academic histories of every author we decided to score each author by their affiliation(s) as reported in the publications, as this would be relatively consistent and thus fairer to all authors in terms of information availability. An archaeology affiliation would have score 1 and a non-archaeology one would have 0. In cases where the affiliation could not be easily established, the website of the institution was examined to look for data on the main research areas, details of the academic position held by the author, or any additional information which could help assign the disciplinary affiliation. In cases where no information could be obtained we erred on the side of caution and assigned a score of 0.5. With these scores, we first calculated an author’s average and used that to obtain the publication’s average which we termed the Archaeology Index.

### Testing the differences in the level of completeness among metadata and its effect size

The Kruskal-Wallis test, a non-parametric test, was used to assess whether there were significant differences in the level of completeness among metadata variables. The post-hoc Dunn test with Bonferroni correction was then used for pairwise comparisons with the R package dunn.test v1.3.6 (Dinno 2024). We computed the effect size of the Kruskal-Wallis tests on the differences between metadata and between publications, respectively, for comparison using the R package rstatix 0.7.2 (Kassambara 2023).

### Correlation analysis and PCA of archaeological metadata

The Spearman correlation was used to measure the strength and direction of the relationships between the metadata variables. The p-values of Spearman’s ρ were adjusted for multiple testing using Bonferroni correction. To further explore and visualize the variation in reporting, PCA was performed using the R package FactoMineR v2.11 (Lê et al. 2008) on the same data (Fig. S6-8).

### Modeling the relationships between metadata completeness and candidate explanatory variables

To model the relationships between metadata completeness and candidate explanatory variables, we applied fractional logistic regression (as called in Stata) (Papke and Wooldridge 1996) which uses a quasi-likelihood approach to handle fractional response variables. This was done via fitting a logistic regression model to estimate the coefficients and estimating robust standard errors using the R package sandwich v3.1.1 (Zeileis et al. 2020) with the ”HC1” estimator (Zeileis 2006). For simple regressions where candidate explanatory variables were modeled separately, the p-values of coefficient estimates were adjusted for multiple testing for each metadata variable using Bonferroni correction. Multiple regressions were performed to address potential confounding between candidate explanatory variables. In these models, the first three PCs instead of the original variables were used due to multicollinearity detected. Finally, the Wald test was used to assess model fit which tests whether at least one coefficient estimate is significantly different from 0.

We also performed extended-support beta (XBX) regression (Cribari-Neto and Zeileis 2010; Kosmidis and Zeileis 2025), which is fully parametric in comparison to the quasi-likelihood based fractional logistic regression. Log-likelihood (log-pseudolikelihood in the case of fractional logistic regression), AIC, BIC and pseudo R^2^ (squared correlation between the predicted and observed values) were calculated to assess goodness of fit (Fig. S20-21). As their pseudo R^2^ were almost identical, we report in the main text the results from the fractional logistic regression model, which is a simpler model that generally resulted in a lower AIC or BIC.

### Assessing multicollinearity in multiple regression analysis

We found significant positive correlations between several pairs of candidate explanatory variables (Fig. S5). This suggests potential multicollinearity in multiple regression which would bias coefficient and standard error estimation. To assess multicollinearity, we computed the variance inflation factor (VIF) for each explanatory variable in each model (Fig. S13-16). A VIF that exceeds 5 or 10 was considered problematic (James et al. 2021).

All analyses were carried out in R v4.4.0 (R Core Team 2018).

## Data Availability

All data used in this study can be found in the Supplementary tables.

## Code Availability

All code used in this study is openly available on GitHub at https://github.com/vicyklee/arch_adna_metadata, under the MIT License.

## Acknowledgments

This research is the result of the synergy project ‘COREX: From Correlations to Explanations: towards a new European prehistory’, funded by the European Research Council (ERC) under the European Union’s Horizon 2020 research and innovation program (Grant Agreement No. 951385). R.S. was additionally supported by the NAWA Polish Returns 2023 project (BPN/PPO/2023/1/00013/U/00001) and the Polish National Science Center Research Component (2024/03/1/HS3/00008). V.Y.K.L. and F.R. were additionally supported by the European Research Council (ERC) under the European Union’s Horizon Europe program (Grant Agreement No. 101077592) and F.R. was also supported by a Novo Nordisk Fonden Data Science Ascending Investigator Award (NNF22OC0076816). We would like to express our thanks to James Fellows Yates for his comments on the pre-print of this paper.

## Author Contributions

R.S., V.Y.K.L., A.T., K.K., S.S., F.R., M.G.T. conceptualized and designed the research; R.S. and V.Y.K.L. collected data; V.Y.K.L., F.R., R.S., M.G.T., A.T. performed data analysis; R.S. and V.Y.K.L. drafted the manuscript; A.T., K.K., S.S., F.R., M.G.T. reviewed, edited and approved the final manuscript.

