## Supplementary Information for "Gaps in archaeological metadata reporting: a meta-analysis of human paleogenomic studies in Western Eurasia"

### Supplementary figures

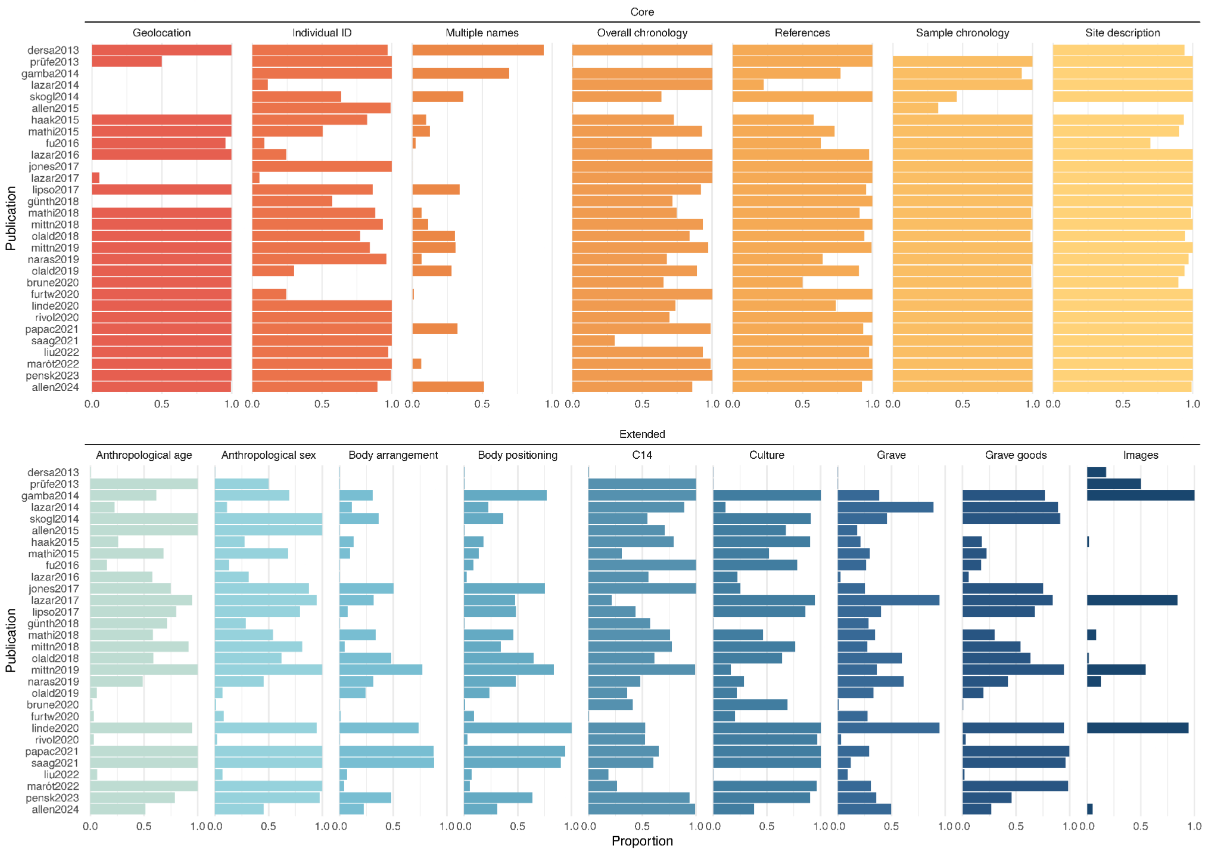


**Figure. S1:** An overview of archaeological metadata reporting in proportions of the 30 peer-reviewed publications from 2013 to 2024.

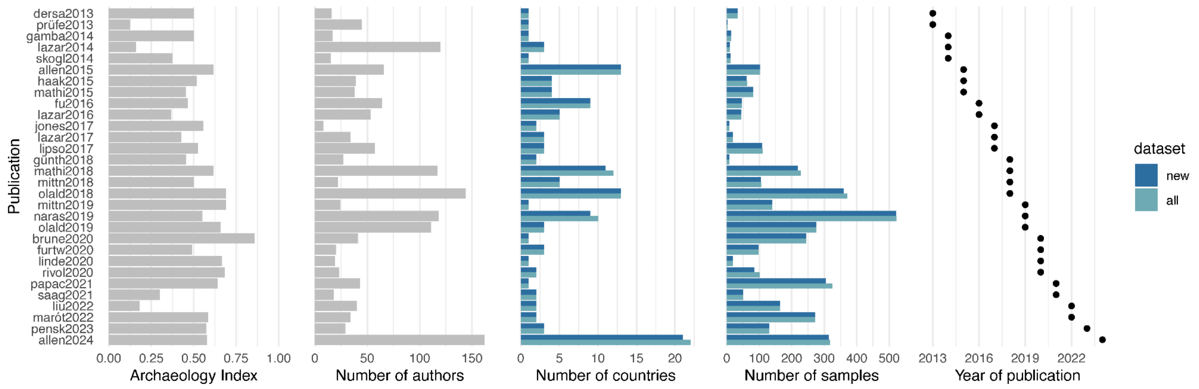


**Figure. S2:** An overview of the candidate explanatory variable including both newly and resequenced sampled individuals for each publication. “New” refers to only newly sampled individuals; ”all” refers to the total dataset.

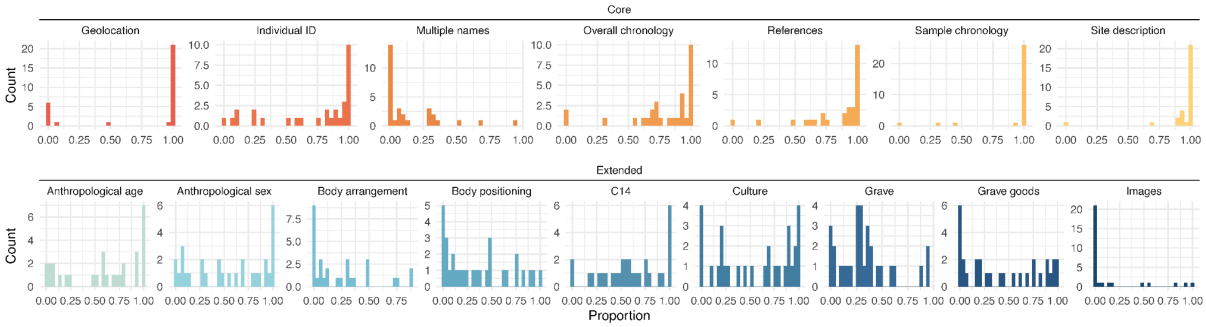


**Figure S3:**. The distributions of metadata completeness.

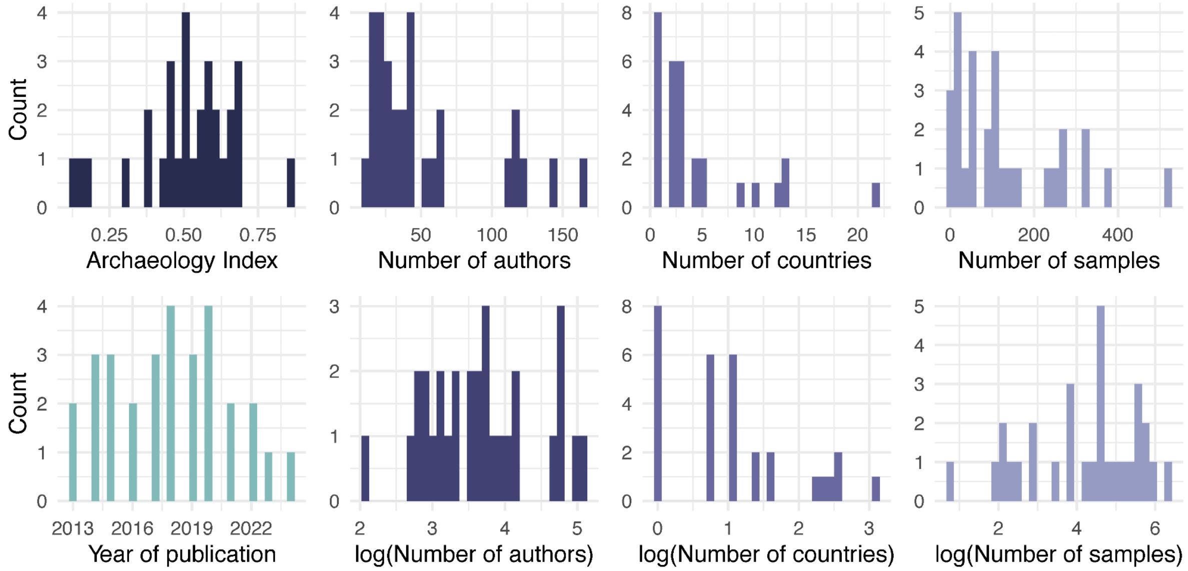


**Figure S4:** The distributions of candidate explanatory variables.

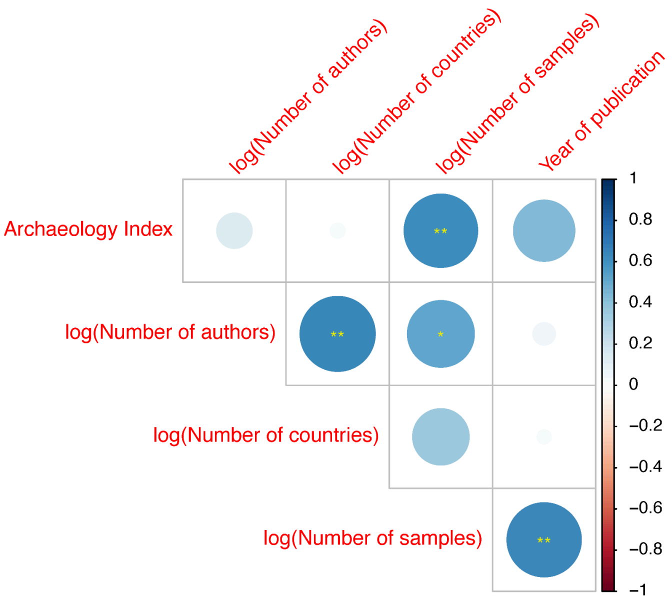


**Figure S5:** Spearman correlations between candidate explanatory variables. Spearman's ρ is indicated by the color and the size of the circle whereas the significance level after Bonferroni correction is indicated by the asterisks ("*": padj < 0.05; "**": padj < 0.01; "***": padj < 0.001).

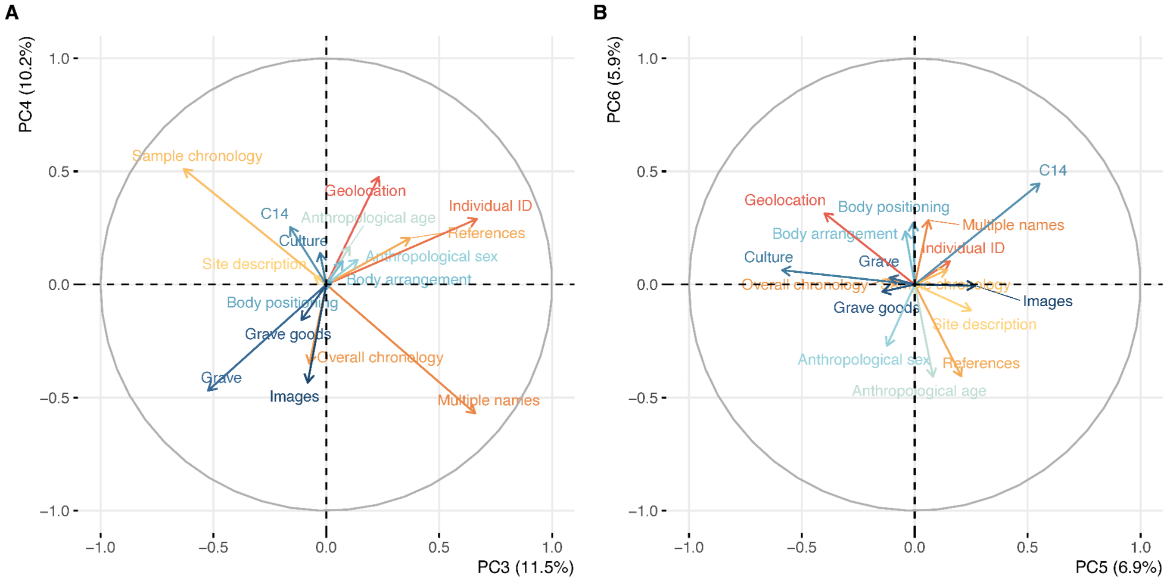


**Figure S6:** The loading plots of PCA. The projected vectors correspond to the contributions of metadata variables to the PCs as measured by correlations. (A) Projections on PC3 and PC4. (B) Projections on PC5 and PC6.

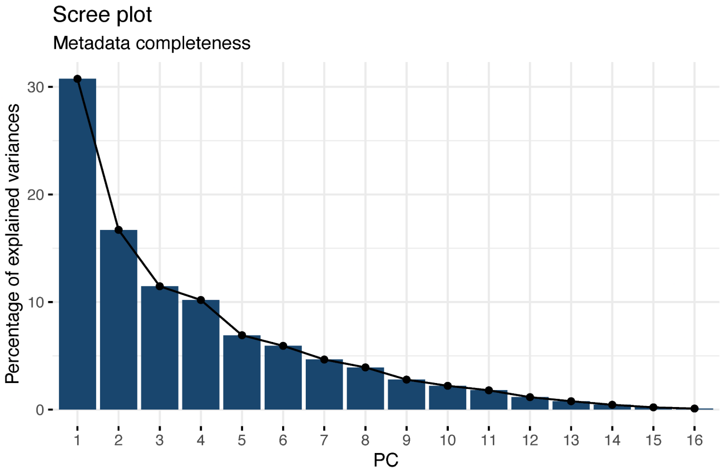


**Figure S7:** The scree plot of PCA of metadata completeness.

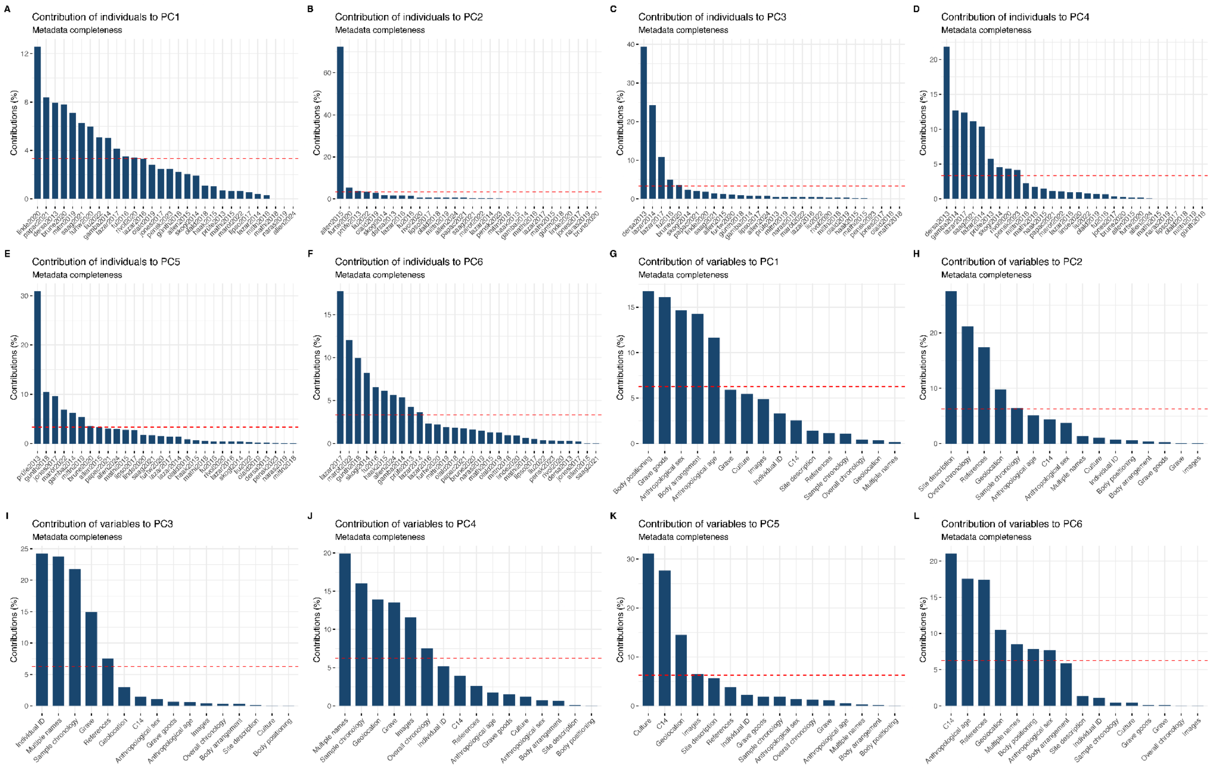


**Figure S8:** Contributions of variables to the first 6 PCs of metadata completeness. (A-F) Contributions of publications. (G-L) Contributions of metadata variables.

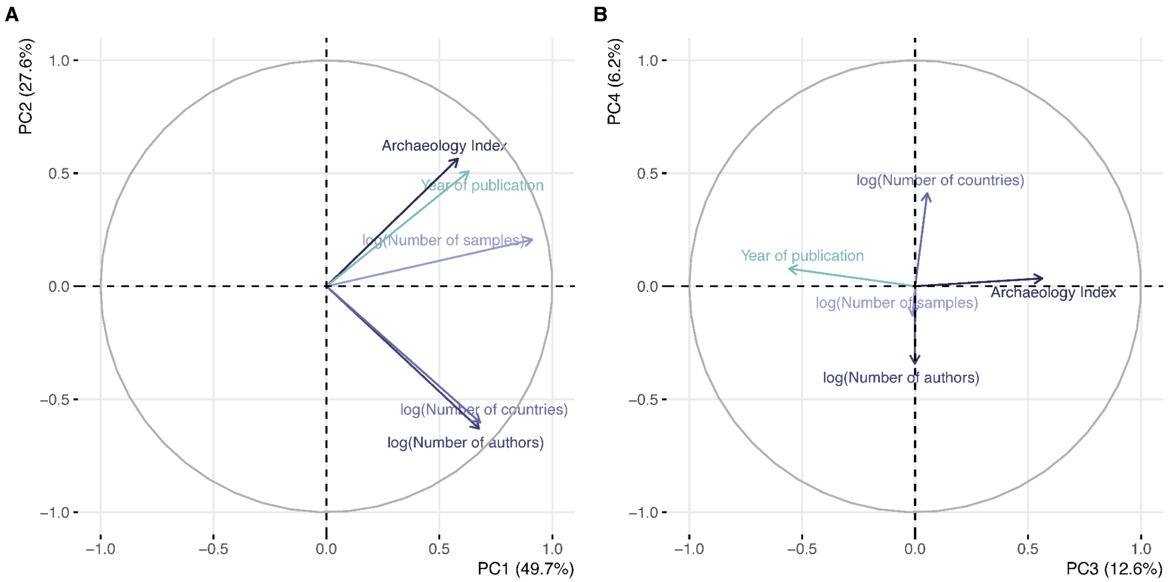


**Figure S9:** The loading plots of PCA. The projected vectors correspond to the contributions of candidate explanatory variables to the PCs as measured by correlations. (A) Projections on PC1 and PC2. (B) Projections on PC3 and PC4.

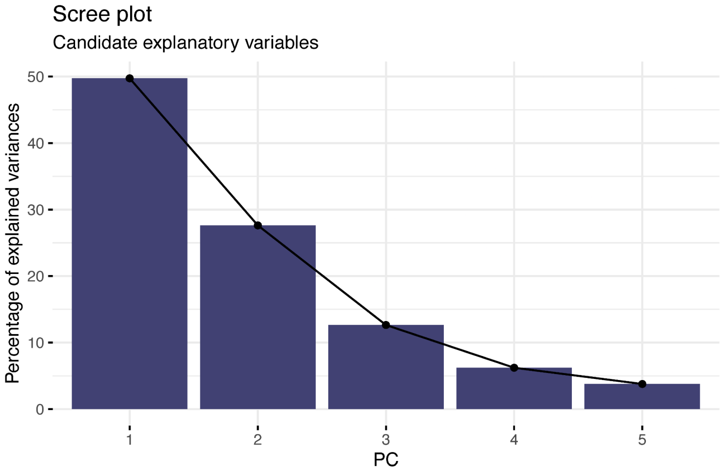


**Figure S10:** The scree plot of PCA of candidate explanatory variables

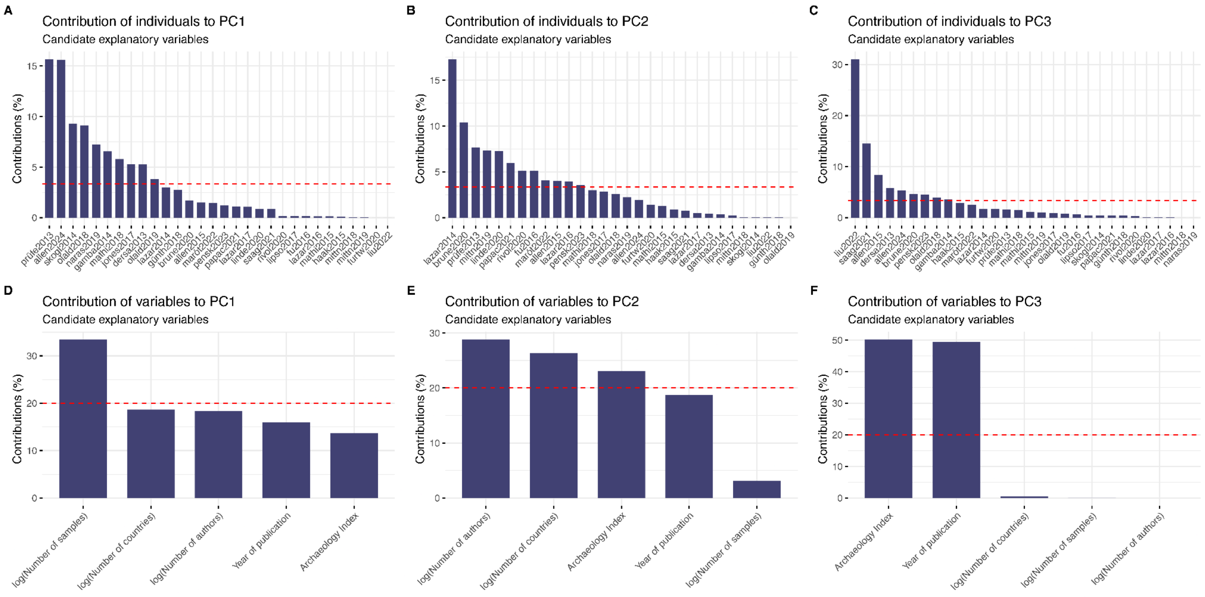


**Figure S11:** Contributions of variables to the first 3 PCs of candidate explanatory variables. (A-C) Contributions of publications. (D-F) Contributions of metadata variables.

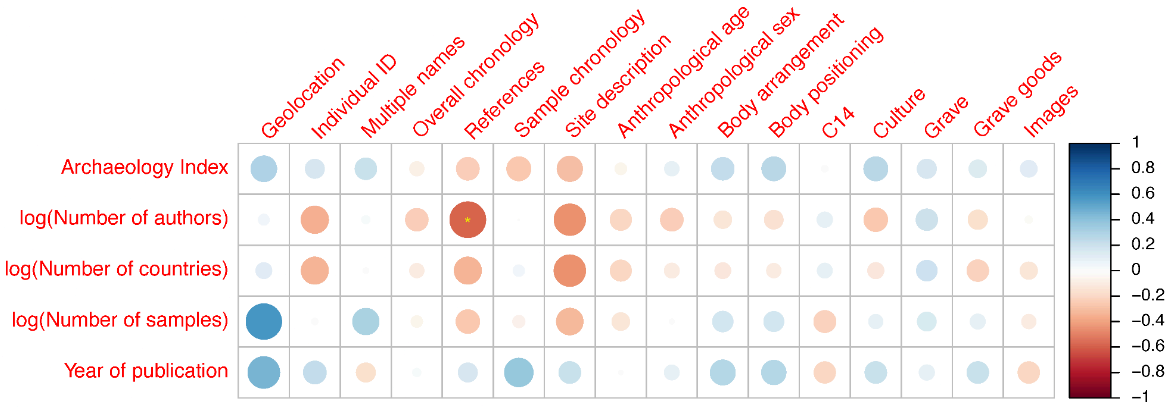


**Figure S12:** Spearman correlations between metadata and candidate explanatory variables. Spearman's ρ is indicated by the color and the size of the circle and the significance level after Bonferroni correction is indicated by the asterisks("*": padj < 0.05; "**": padj < 0.01; "***": padj < 0.001).

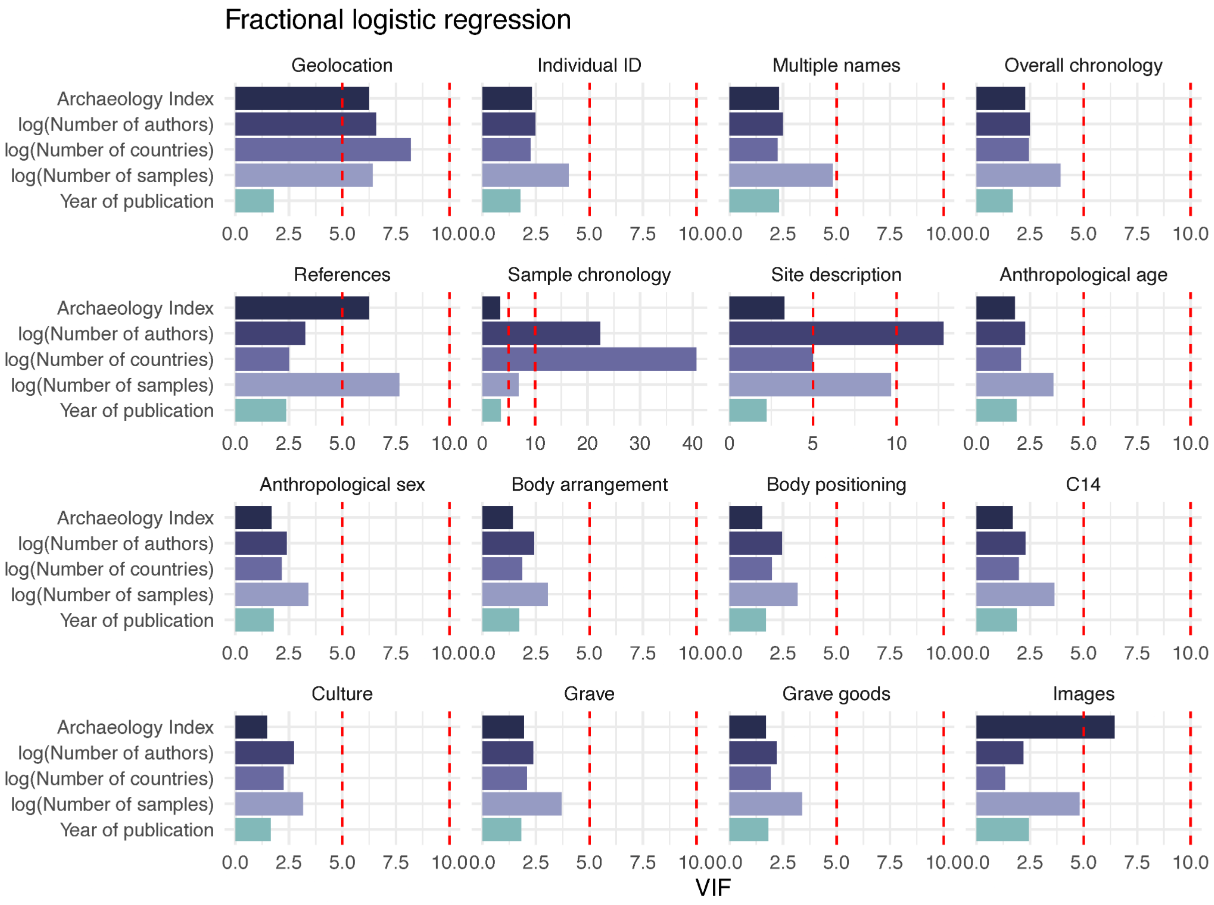


**Figure S13:** VIFs of fractional logistic regression models.

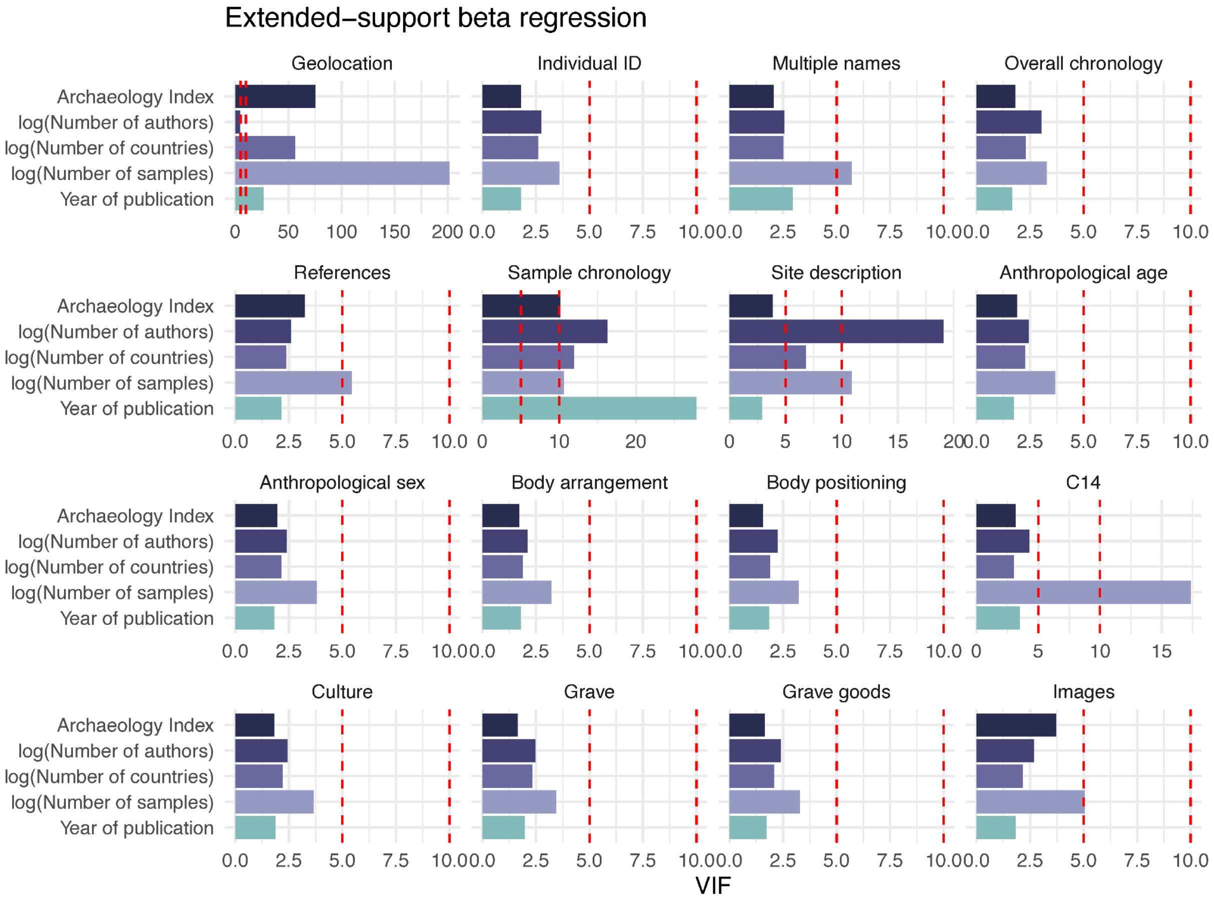


**Figure S14:** VIFs of XBX regression model.

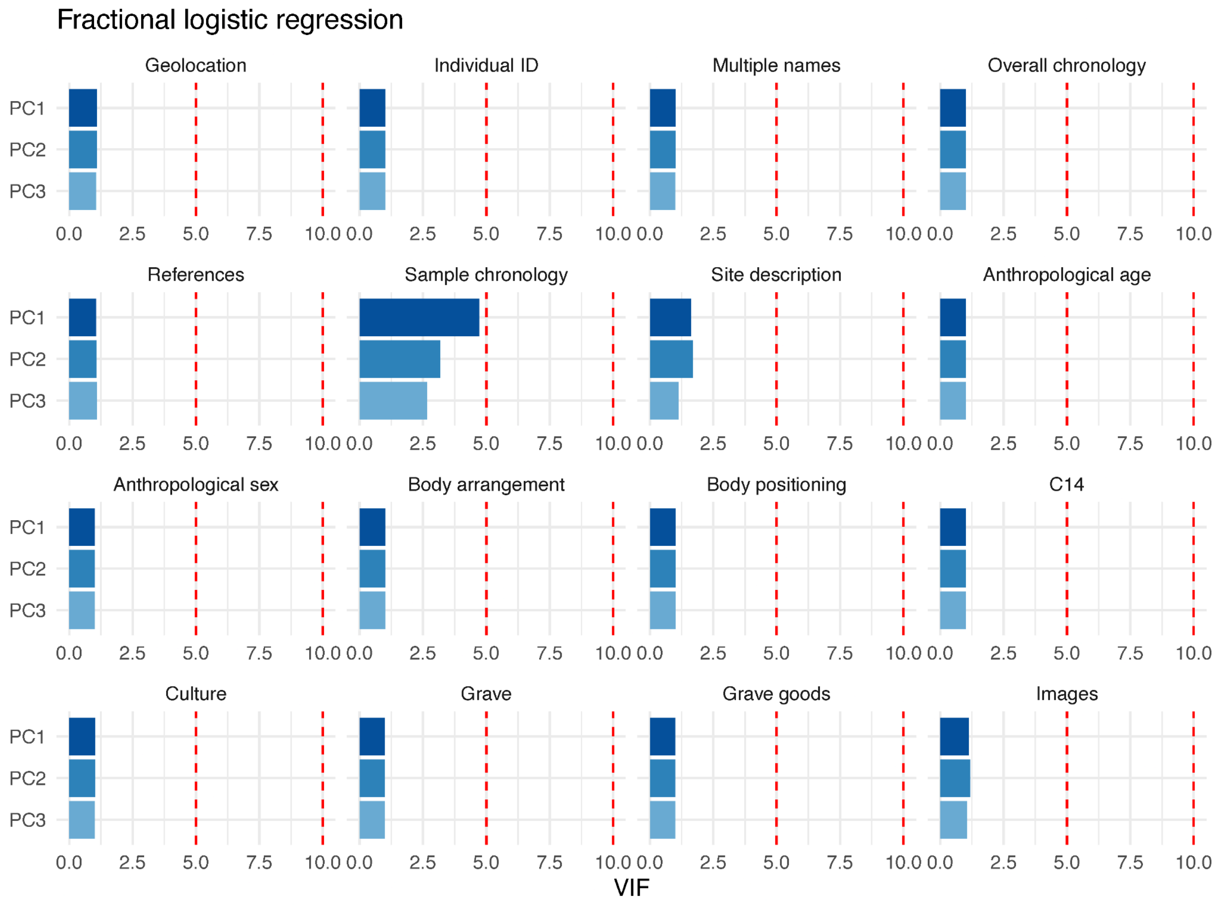


**Figure S15:** VIFs of fractional logistic regression models with the first three PCs of the original variables as regressors.

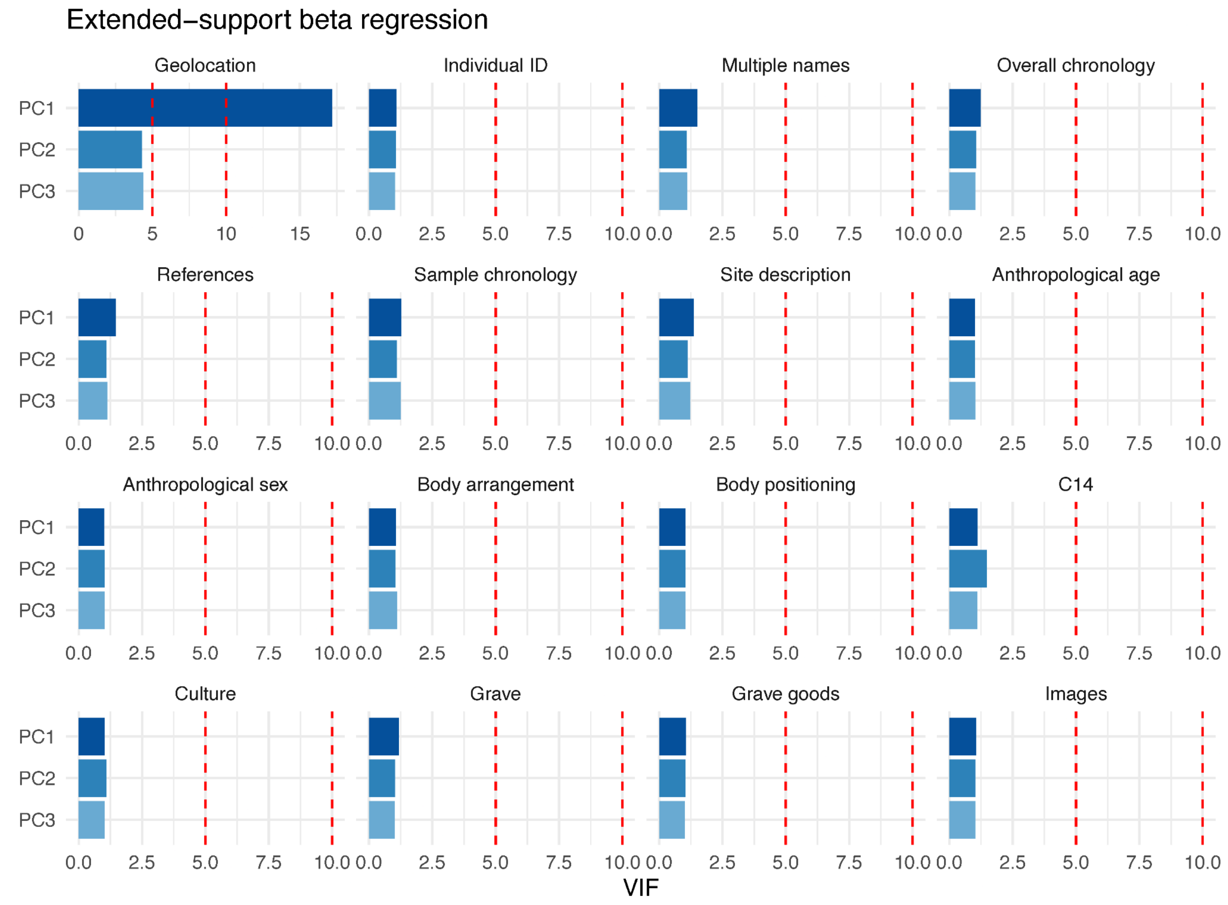


**Figure S16:** VIFs of XBX regression models with the first three PCs of the original variables as regressors.

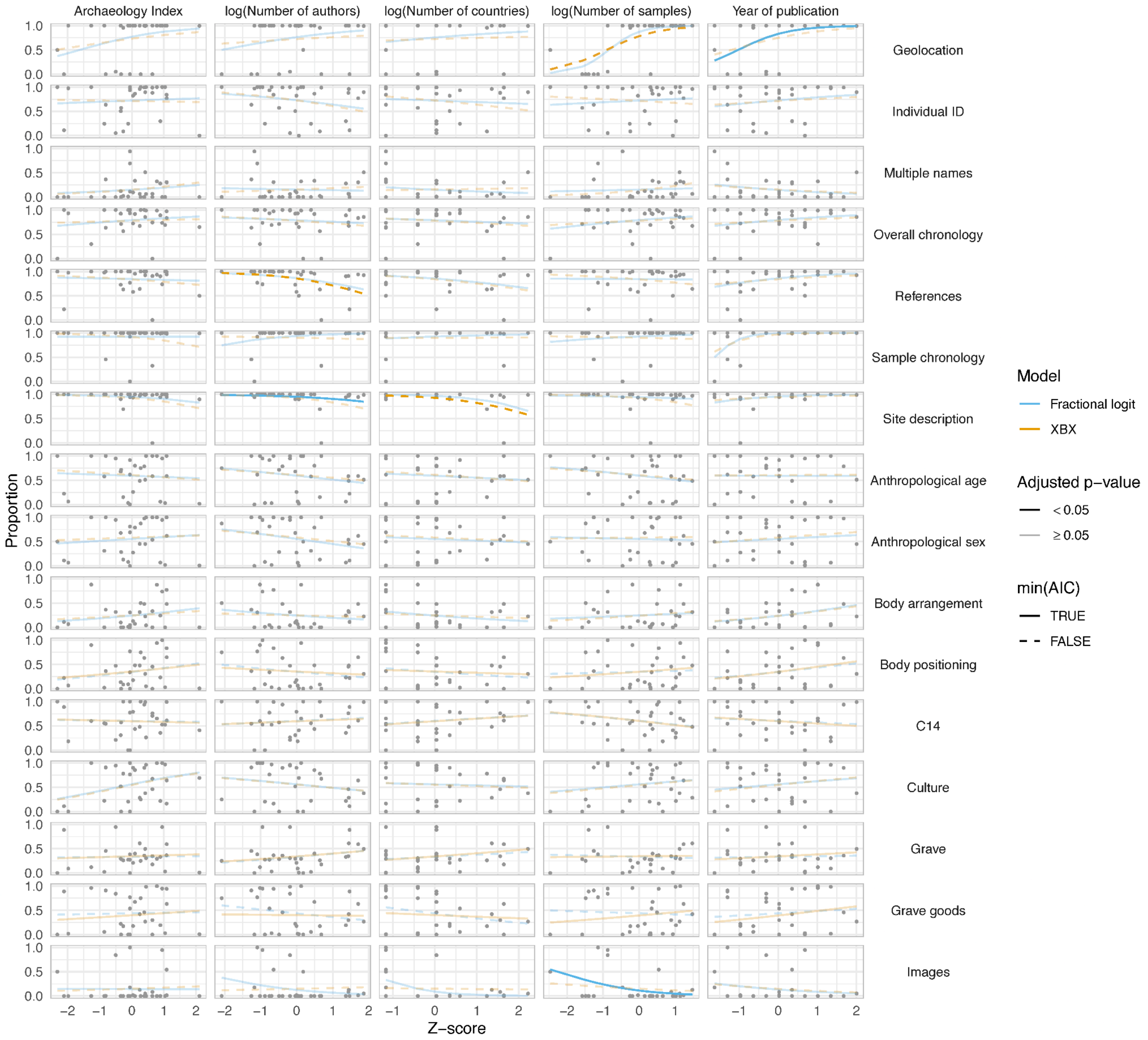


**Figure S17:** Model fit of fractional logistic and XBX regression models. An opaque line indicates the coefficient estimate ($\hat{\beta}$) is significant after Bonferroni correction; a solid line indicates the model with the lower AIC.

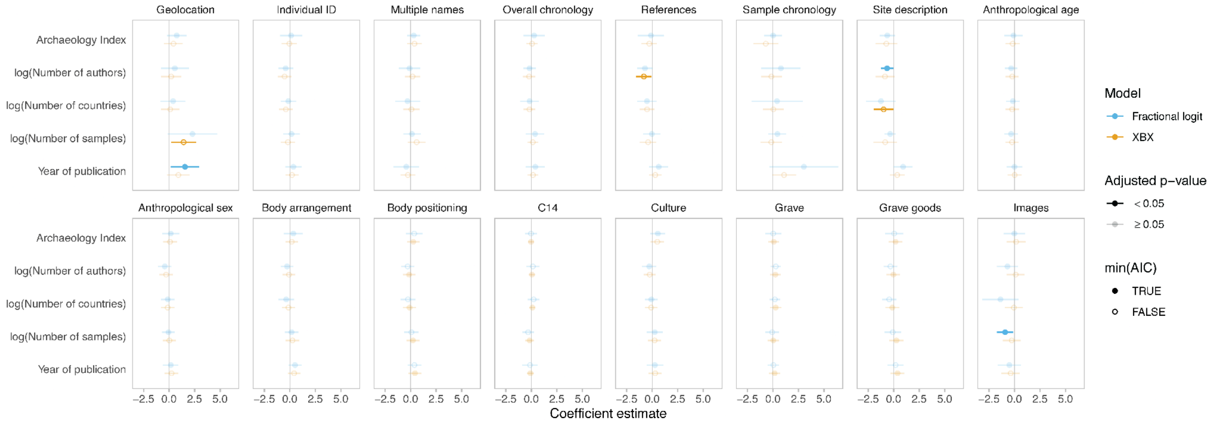


**Figure S18:** The effects of candidate explanatory variables on metadata completeness. The line represents the 95% CI around the coefficient estimate (β̂). An opaque line indicates the coefficient estimate is significant after Bonferroni correction; a solid circle indicates the model with the lower AIC.

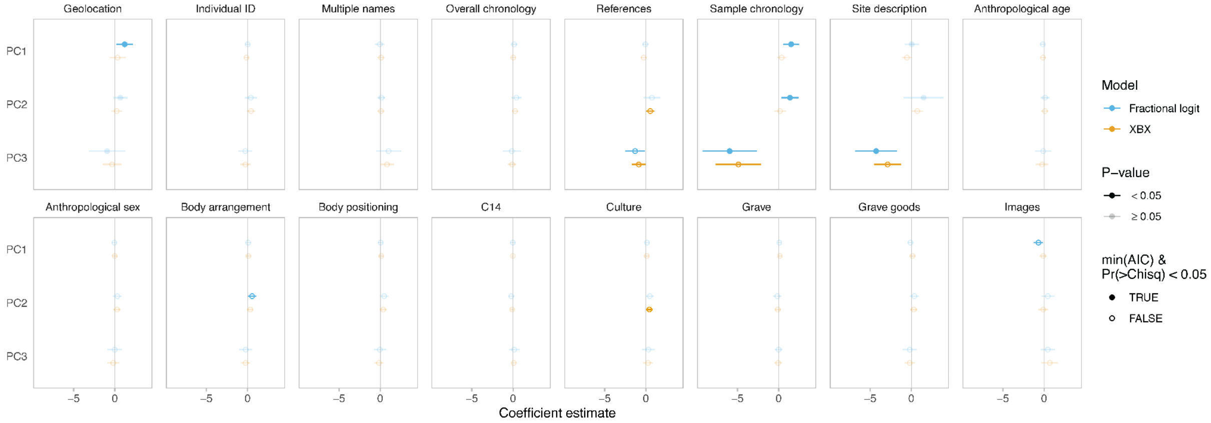


**Figure S19:** The effects of the first three PCs of the original variables on metadata completeness. The line represents the 95% CI around the coefficient estimate. An opaque line indicates the coefficient estimate is significant; a solid circle indicates the model with the lower AIC that passes the Wald test for model fit.

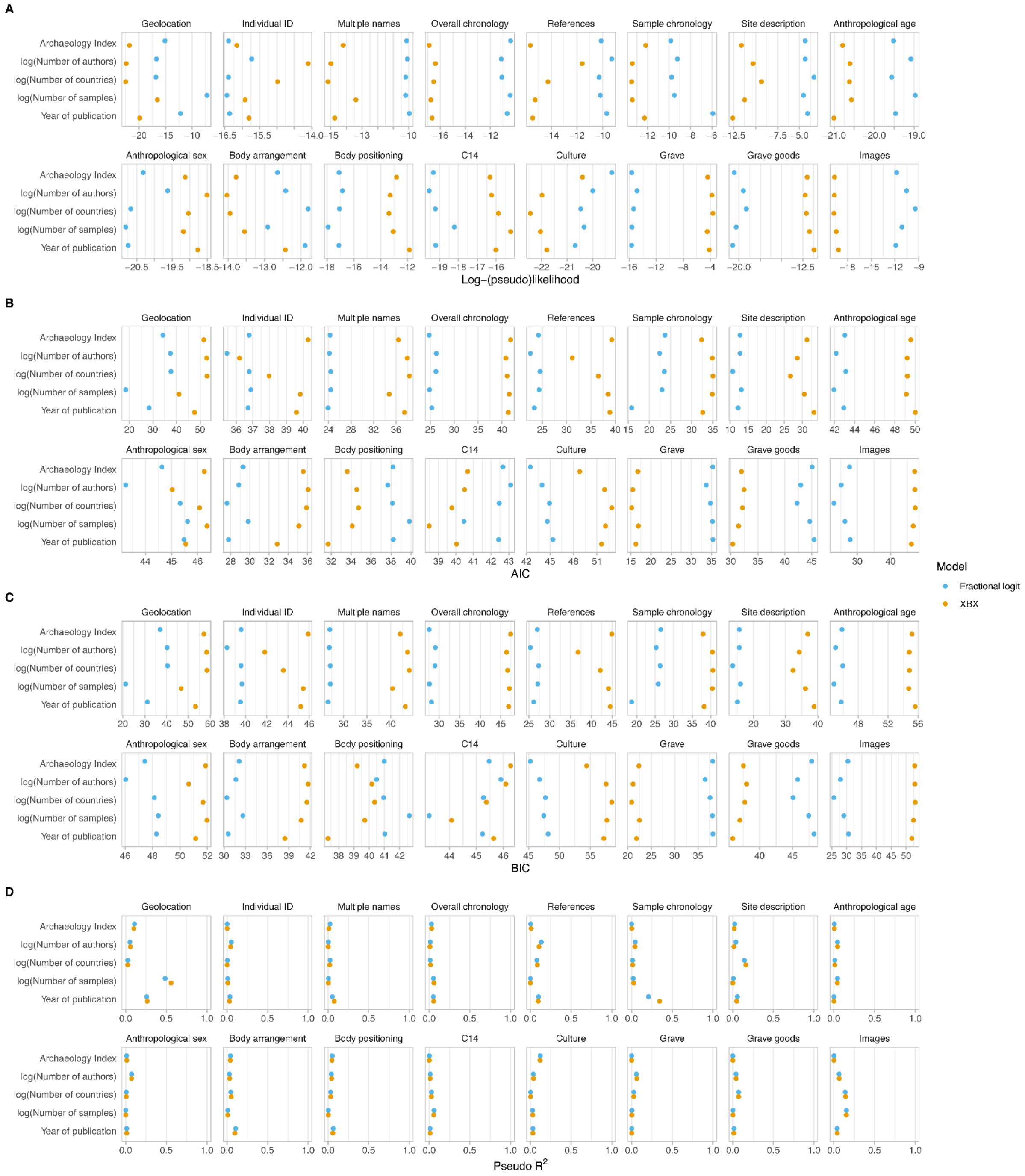


**Figure S20:** Model comparisons between fractional logistic and XBX regression models in simple regressions. (A) Log-(pseudo*)likelihood. (B) AIC. (C) BIC. (D) Pseudo R2, computed as the squared correlation of the predicted and observed values. *Log-pseudolikelihood applies to the fractional logistic regression model.

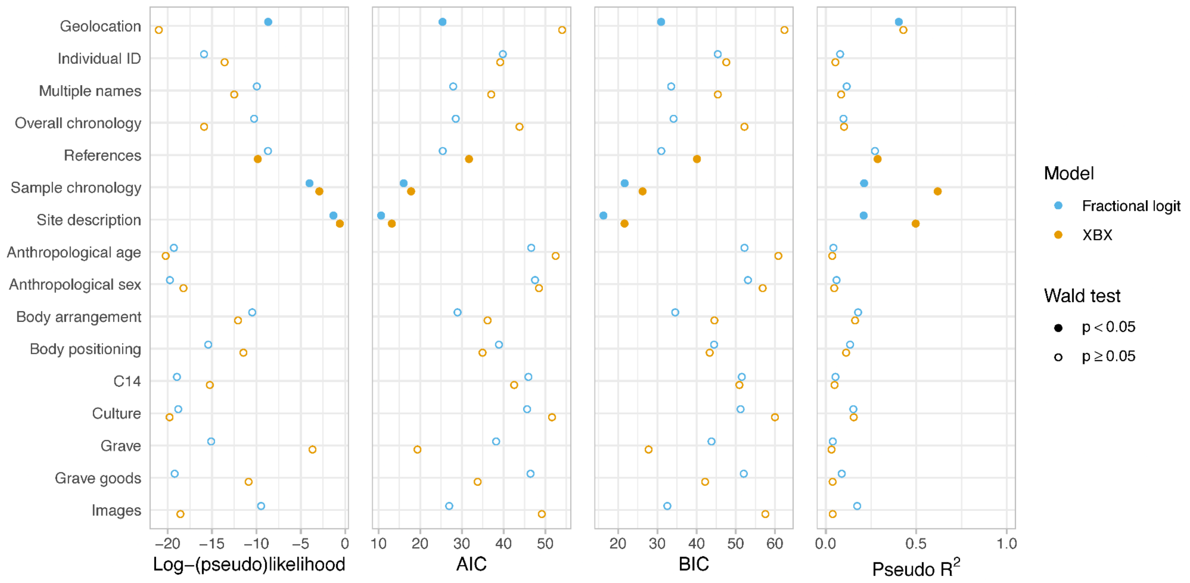


**Figure S21:** Model comparisons between fractional logistic and XBX regression models in multiple regressions. Log-pseudolikelihood applies to the fractional logistic regression model. Pseudo $R^2$ is computed as the squared correlation of the predicted and observed values.

### Supplementary tables (separate file)

**Table S1:** Included in the file as spreadsheet "Table S1". Summary of bibliographic data of papers used for the study.

**Table S2:** Included in the file as spreadsheet "Table S2". Summary of institutional affiliations reported in the sampled studies and their classification along the genetic-archaeological spectrum.

**Table S3:** Included in the file as spreadsheet "Table S3". Summary of cited author's institutional affiliations reported in the sampled studies.

**Table S4:** Included in the file as spreadsheet "Table S4". Summary of metadata survey per reported individual.

**Table S5:** Included in the file as spreadsheet "Table S5". Encoding table.

**Table S6:** Included in the file as spreadsheet "Table S6". Dunn's Kruskal-Wallis multiple comparisons (p-values adjusted with the Bonferroni method).

**Table S7:** Included in the file as spreadsheet "Table S7". Preliminary template.
